# *Vibrio cholerae* Type VI Secretion System Auxiliary Cluster 3 is a Pandemic-associated Mobile Genetic Element

**DOI:** 10.1101/868539

**Authors:** Francis J. Santoriello, Lina Michel, Daniel Unterweger, Stefan Pukatzki

## Abstract

All sequenced *Vibrio cholerae* isolates encode a contact-dependent type VI secretion system (T6SS) in three loci that terminate in a toxic effector and cognate immunity protein (E/I) pair, allowing for competitor killing and clonal expansion in aquatic environments and the host gut. Recent studies have demonstrated variability in the toxic effectors produced by different *V. cholerae* strains and the propensity for effector genes to undergo horizontal gene transfer. Here we demonstrate that a fourth cluster, auxiliary cluster 3 (Aux3), encoding the E/I pair *tseH/tsiH*, is located directly downstream from two putative recombinases and is flanked by repeat elements resembling *att* sites. Genomic analysis of 749 *V. cholerae* isolates, including both pandemic and environmental strains, revealed that Aux3 exists in two states: a ∼40 kb prophage-like element in nine environmental isolates and a ∼6 kb element in pandemic isolates. These findings indicate that Aux3 in pandemic *V. cholerae* is evolutionarily related to an environmental prophage-like element. In both states, Aux3 excises from the chromosome via site-specific recombination to form a circular product, likely priming the module for horizontal transfer. Finally, we show that Aux3 can integrate into the Aux3-naïve chromosome in an integrase-dependent, site-specific manner. This highlights the potential of Aux3 to undergo horizontal transfer by a phage-like mechanism, which based on pandemic coincidence may confer currently unknown fitness advantages to the recipient *V. cholerae* cell.

**Significance Statement:** *V. cholerae* is a human pathogen that causes pandemics affecting 2.8 million people annually (1). The O1 El Tor lineage is responsible for the current pandemic. A subset of non-O1 strains cause cholera-like disease by producing the major virulence factors cholera toxin and toxin co-regulated pilus but fail to cause pandemics. The full set of *V. cholerae* pandemic factors is unknown. Here we describe the type VI secretion system (T6SS) Aux3 element as a largely pandemic-specific factor that is evolutionarily related to an environmental prophage-like element circulating in non-pathogenic strains. These findings shed light on *V. cholerae* T6SS evolution and indicate the Aux3 element as a pandemic-enriched mobile genetic element.

## Introduction

*Vibrio cholerae*, the causative agent of cholera, is capable of causing natural pandemics. O1 Classical strains caused the first six pandemics, and O1 El Tor strains cause the current 7^th^ pandemic (2–4). Pandemic strains cause diarrheal disease with the major virulence factors cholera toxin (CT) and toxin co-regulated pilus (TCP) (5–8). Several non-O1 strains, however, carry these main virulence factors and cause isolated cases of cholera-like illness without causing pandemic outbreaks (9–11). The full set of factors that make a given *V. cholerae* strain pandemic is unknown.

In its aquatic reservoir and the human small intestine, *V. cholerae* competes with other bacteria and predatory eukaryotic cells. *V. cholerae* employs the type VI secretion system (T6SS), a contractile nanomachine resembling a T4 bacteriophage tail that kills neighboring competitors through the contact-dependent translocation of toxic effector proteins (12–14). The main components of the T6SS are encoded in three loci (the large cluster, auxiliary cluster 1 (Aux1), and auxiliary cluster 2 (Aux2)), each terminating in an effector/immunity (E/I) pair (13, 15, 16). While T6 effectors are toxic to distinct bacteria, kin cells are protected by cognate immunity proteins (16, 17). It is hypothesized that this allows a strain to clear a niche and propagate clonally (18). Comparative genomic studies of *V. cholerae* T6SS loci have demonstrated that all pandemic strains carry an identical set of effector genes referred to as A-type effectors but environmental strains encode variable effector and immunity subtypes (19, 20). Pan-genome hierarchical clustering of *V. cholerae* does not reflect the dispersion of these effector subtypes (20), suggesting horizontal gene transfer (HGT) as a critical player in T6SS E/I evolution. *V. cholerae* in both the estuarine environment and its human host is exposed to exogenous DNA, bacteriophage, and conjugative elements. Further, chitin-associated *V. cholerae* upregulates the T6SS and natural competence machinery (21–23), driving rapid evolution via inter- and intra-species competition. Recently, horizontal transfer of variable T6SS effectors to chitin-bound *V. cholerae* was demonstrated *in vitro* (24). These studies indicate the aquatic environment and potentially the host gut as a reservoir for the acquisition of new E/I subtypes.

Some T6SS components are bacteriophage structural homologs (25–27), suggesting that the T6SS is the repurposing of one or more lysogenic phage infections. *V. cholerae* T6SS loci do not, however, reflect typical prophage genomic organization or encode functional recombinases. Several other virulence factors in *V. cholerae* such as CT, TCP, and the SXT element are either phage-encoded or acquired by phage-like site-specific recombination (28–32). In each case, integrases and transposases catalyze recombination. These data support the idea that bacteriophage and phage-derived elements play an integral role in the development of pandemic *V. cholerae* strains.

Recently, Altindis *et al*. identified a fourth T6SS cluster in *V. cholerae*. This locus, designated Auxiliary cluster 3 (Aux3), encodes a PAAR adaptor protein, a hydrolase (TseH), and its cognate immunity protein (TsiH) (33). TseH is likely loaded onto the tip of the T6SS with the assistance of the PAAR adaptor. After translocation into the target cell, TseH catalyzes peptidoglycan degradation. Periplasmic TsiH expression neutralizes this degradation. Unlike the three core T6SS loci, Aux3 is not conserved in all sequenced *V. cholerae* strains (34). Here, we demonstrate that Aux3 extends upstream to include an integrase and a transposase, and that phage-like *att* sites flank the region from the integrase to *tsiH*. By analyzing 749 *V. cholerae* genomes, we determined that the Aux3 element is found in 572 strains of which 566 encode CT, TCP, and a pandemic A-type T6SS effector set (19, 20). Based on phylogenetic analysis of a subset of strains, we show that the Aux3 locus appears to have expanded within the entire pandemic lineage. We further determined that Aux3 is present in nine non-pandemic environmental isolates. The environmental Aux3 locus, however, encodes 42-47 extra bacteriophage homologs and appears to move by HGT, indicating that the pandemic Aux3 locus is likely the evolutionary remnant of a prophage-like element circulating in the aquatic reservoir. Finally, both the environmental and pandemic versions of Aux3 excise from the genome by site-specific recombination, but only the environmental Aux3 integrase can catalyze transfer to a naïve strain. These findings highlight this locus as a pandemic-associated mobile genetic element (MGE).

## Results

### Phage-like *att* Sites Flank T6SS Cluster Aux3

Analysis of the Aux3 locus in O1 El Tor strain N16961 revealed that the genes encoding *PAAR, tseH*, and *tsiH* (VCA0284-VCA0286) are immediately downstream from two genes annotated as “phage integrase” (VCA0281, *int*) and “IS5 transposase” (VCA0282, *is5*) (Fig. 1*A*). Sliding-window analysis of the region from VCA0280-VCA0287 reveals blocks of variable GC content within Aux3 compared to the surrounding genomic flanks (Fig. 1*A*). Based on this proximity to putative recombinases and the differential GC content of this region, we hypothesized that this locus constitutes a potential MGE. Recombinase-encoding MGEs are often flanked by repeat elements (attachment (*att*) sites) that serve as the locus of enzyme binding and DNA recombination. Alignment of Aux3-encoding and Aux3-naïve *V. cholerae* strains reveals a single recombinant site on either side of Aux3 indicative of site-specific recombination (*SI Appendix*, Fig. S1*A*). We thus probed the intergenic sequences between *gcvT* and *int* as well as *tsiH* and *thrS* for repetitive sequences and identified two long, direct repeats (referred to as *attL1/attL2* and *attR1/attR2*) separated by approximately 40 bp on either side of the Aux3 locus (Fig. 1*B*), the second of which contains a stretch of six thymidines resembling the λ phage *att* site (35). Alignment of the *attL1/attL2* and *attR1/attR2* sequences from O1 El Tor strain N16961 with *attC1/attC2* from environmental strain DL4215 shows strong homology upstream of *attL2/attC2* and downstream of *attR2/attC2*. These results indicate that the sequence between *attL2* and *attR2* is Aux3-derived, while the sequence outside these sites is derived from the Aux3-naïve genome. We propose that *att2* is the relevant *att* site for Aux3 recombination (*SI Appendix*, Fig. S1 *B* and *C*). Importantly, *attL2* and *attR2* are found flanking the Aux3 cluster in all analyzed Aux3-encoding *V. cholerae* strains, and *attC2* exists in a single copy between *gcvT* and *thrS* in all Aux3-naïve strains (Fig. 1*C*). These findings demonstrate that the Aux3 cluster extends from VCA0281-VCA0286 and potentially constitutes a MGE capable of excising from the genome by site-specific recombination.

**Figure 1.**
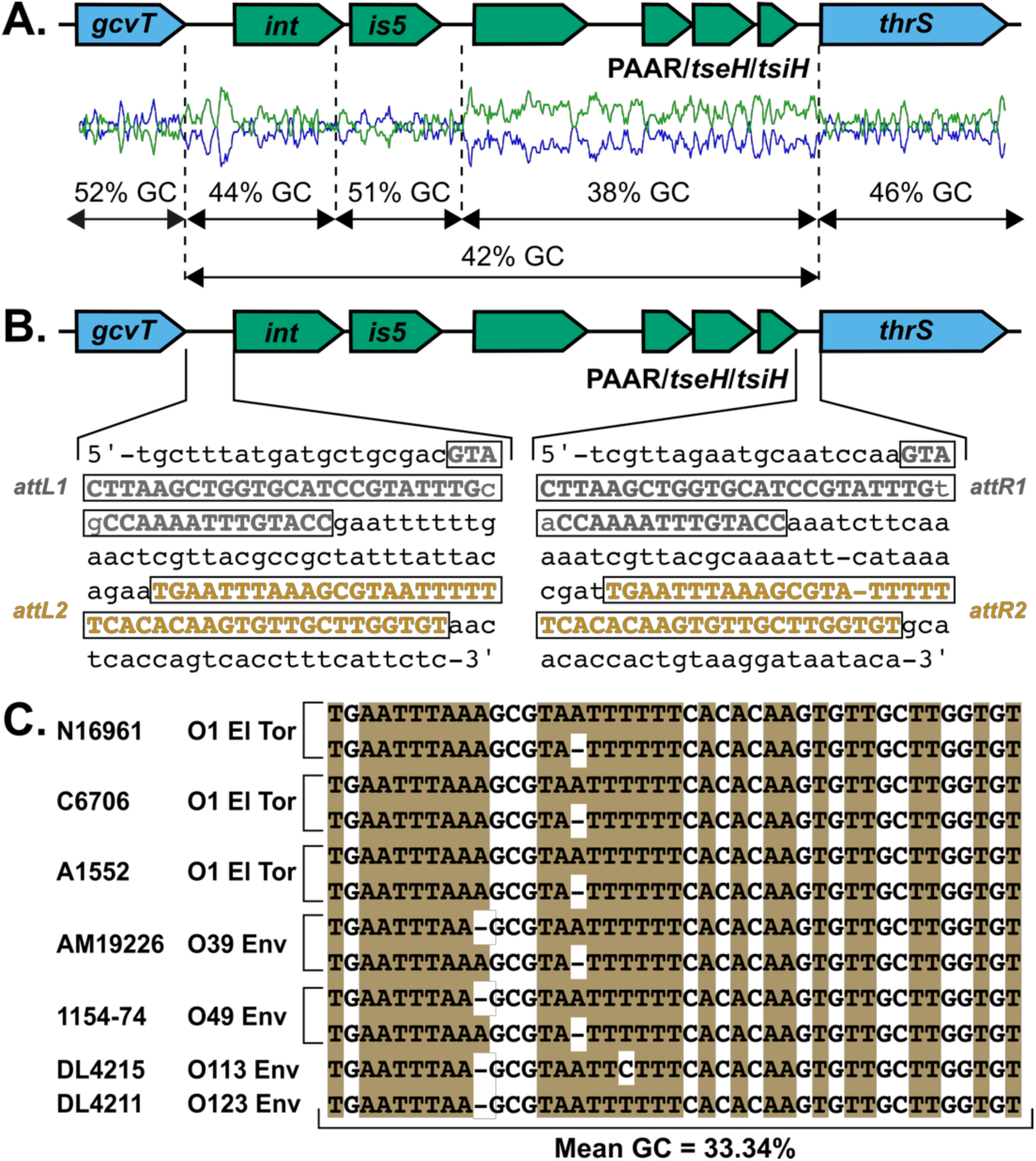
T6SS Aux3 module is flanked by conserved phage-like *att* sites. (*A*) Local GC content was analyzed for the N16961 Aux3 cluster and the flanking regions. Aux3 genes are shown in green and the genomic flanks in blue. GC content (blue line) and AT content (green line) were generated using a sliding window of 40 bp. (*B*) Intergenic regions flanking Aux3 were analyzed for repeated sequences. Direct repeat sequences are boxed and shown in grey (*attL1/attR1*) and orange (*attL2/attR2*). (*C*) Alignment of phage-like *att* site *att2* from Aux3-encoding strains (N16961, C6706, A1552, AM-19226, 1154-74) and Aux3-naïve environmental strains (DL4215, DL4211). Both *attL2* (top) and *attR2* (bottom) are represented for Aux3-encoding strains, and *attC2* is shown for each naïve strain. The average GC content is shown, as *att* sites are typically AT-rich regions.

### Aux3 is Conserved and Enriched in O1 Pandemic Strains

A BLASTN search of the El Tor N16961 Aux3 module in 25 pandemic and environmental *V. cholerae* genomes revealed strong conservation of the Aux3 module in pandemic O1 strains of both the Classical and El Tor biotypes (*SI Appendix*, Fig. S1*A*). All analyzed environmental strains lacked the Aux3 cluster (*SI Appendix*, Fig. S1*A*). To determine the scale of Aux3 enrichment in pandemic *V. cholerae* strains, we probed the coincidence of *tseH* with the pandemic A-type T6SS effectors *tseL, vasX*, and *vgrG3* (19, 20), as well as *ctxAB* and *tcpA*. We performed a Megablast search for these six loci across 749 *V. cholerae* genomes from the PATRIC database (36) to determine the grade (a weighted score accounting for query coverage as well as pairwise identity) for each locus in each genome. Strains were grouped based on having > 99% grade to *tseH* as well as > 99% grade to the A-type effectors. Of a total 547 strains with hits for *tseH*, 461 strains had a grade of > 99% for *tseH, tseL, vasX*, and *vgrG3*, corresponding to an enrichment of *tseH* in pandemic strains of p = 2.2×10^−16^ by Fisher’s Exact Test (*SI Appendix*, Table S4). Due to the fragmented nature of available *V. cholerae* genomes in the PATRIC database, this enrichment is likely an underestimation. We expanded our analysis to include all strains with *tseH* regardless of grade and found that 566 of 572 *tseH*-encoding strains also encoded all five selected pandemic factors (*SI Appendix*, Fig. S2*A*). It is important to note that the Aux3 element is absent from non-O1/O139 pathogenic strains that do not cause pandemics but carry the major virulence factors CT and TCP (*SI Appendix*, Fig. S3). These data demonstrate that Aux3 is enriched in the subset of *V. cholerae* strains with the largest global impact on health.

### The Pandemic Aux3 Module Is Evolutionarily Related to a Prophage-like Element

Our Megablast search for *tseH, tseL, vasX*, and *vgrG3* in the *V. cholerae* genomes in the PATRIC database revealed six *tseH-*encoding strains that lack *tseL* and *ctxAB* (*SI Appendix*, Fig. S2 *A* and *B*). Three of these strains are environmental O1 strains (2012Env-9, Env390, and 2479-86), two of which encode the toxin co-regulated pilus (2012Env-9 and Env390). The remaining three strains (AM-19226, 1154-74, and P-18748) are nonO1/O139 isolates. Investigation of the region between *gcvT* and *thrS* in these strains revealed an Aux3 locus approximately 40kb in length compared to the 6kb-long module found in pandemic strains (Fig. 2*A*). A Megablast search for this region in NCBI returned three more strains with this elongated Aux3 element (*V. cholerae* str. 20000, *Vibrio sp*. 2015V-1076, and *Vibrio sp*. 2017V-1038). Importantly, *attL2* and *attR2* flank the Aux3 region in each of these strains (Fig. 1*C*). Alignment of the Aux3 region in these nine environmental strains reveals variability in the additional sequence between VCA0281 and VCA0283, with most of the variability in the 5’ half of the region. Further, all environmental strains lack VCA0282 (*SI Appendix*, Fig. S4). Analysis of these nine environmental strains by PHASTER (37) predicts that the Aux3 region in non-pandemic strains resembles an intact prophage in both gene content and organization (Fig. 2*B* and *SI Appendix*, Fig. S5). These data support the idea that Aux3 exists in two basic states, environmental Aux3 (Aux3^E^) and pandemic Aux3 (Aux3^P^), with a common prophage origin. We performed a core genome alignment of 69 pandemic and environmental *V. cholerae* strains as well as 8 *Vibrio sp*. and one *V. mimicus* isolate, which shows that the incidence of Aux3 in environmental strains is not reflective of phylogeny (Fig. 3). This scenario leads us to conclude that while Aux3^P^ likely expanded clonally in pandemic strains, Aux3^E^ may circulate environmentally by HGT. We hypothesize that the evolution of Aux3^P^ in the pandemic lineage began with the integration of a horizontally transferred phage-like element which then underwent a large deletion event to generate the smaller module (Fig. 2*C*). We cannot, however, rule out the inverse event, in which Aux3^P^ gained excess prophage-related genes in a large insertion event to form Aux3^E^. As all Aux3^E^ strains lack *is5* (VCA0282) (*SI Appendix*, Fig. S4), leading us to assume that the insertion of this element occurred in an evolutionary intermediate (Fig. 2*C*). We have not yet identified a strain encoding this intermediate Aux3^P^ that lacks the IS5 element.

**Figure 2.**
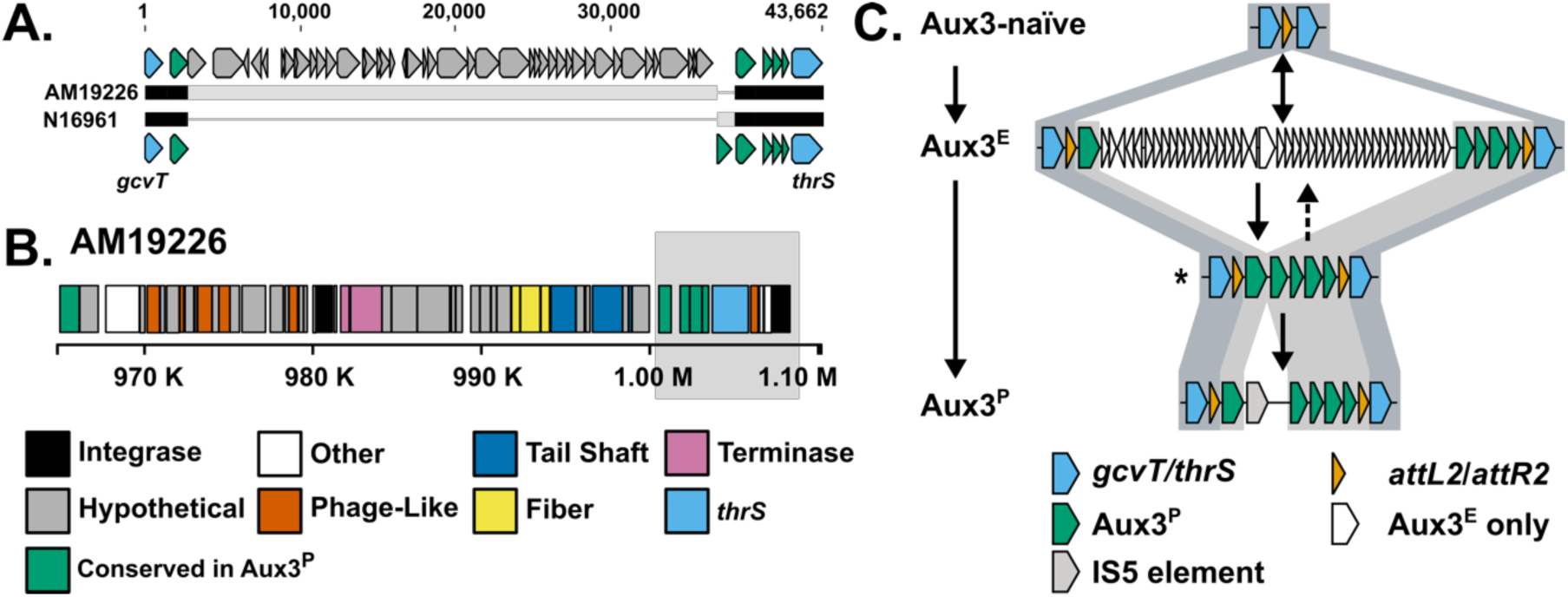
Environmental *tseH*-encoding *V. cholerae* strains carry an alternative prophage-like Aux3 element. (*A*) MAUVE alignment of the Aux3 locus from pandemic strain N16961 and environmental strain AM-19226. Flanking genes are shown in blue. Pandemic Aux3 genes are shown in green. Black bars indicate nucleotide agreements and grey bars indicate differences. (*B*) PHASTER genome diagram showing predicted Aux3 prophage region from the leading integrase (VCA0281) through the superintegron integrase (VCA0291) in AM-19226. Coding regions are colored according to homology to broad categories of known phage genes. Light grey box indicates region not called by PHASTER, but verified manually. (*C*) Schematic of proposed Aux3 module evolution from Aux3-naïve environmental to Aux3^P^ strains. Conserved regions between steps are highlighted in light blue (environmental to pandemic Aux3^P^) or grey (Aux3^E^ to Aux3^P^). * indicates a putative unseen intermediate stage in Aux3 evolution. Dashed arrow indicates alternate hypothesis of a large insertion to form Aux3^E^.

**Figure 3.**
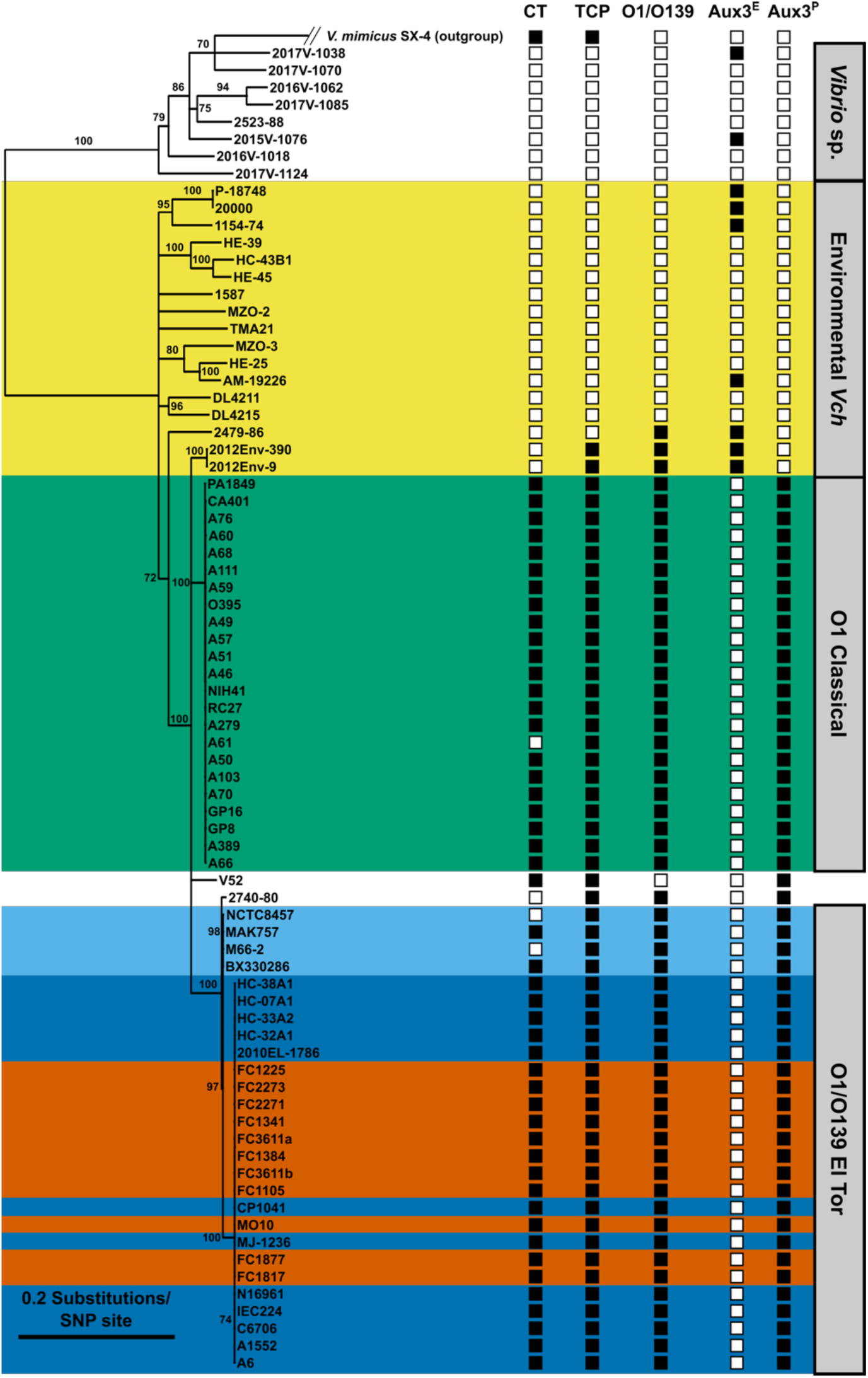
The Aux3 element is enriched in pandemic *V. cholerae* and sporadically distributed in environmental strains. A phylogenetic tree was constructed using the GTR + Gamma Maximum likelihood model in RAxML based on core genome SNP alignment of 69 *V. cholerae*, 8 *Vibrio* sp., and 1 *V. mimicus* genome sequences. Bootstrapping support values are indicated next to their respective branches. Nodes with support values < 70 were collapsed. Presence (black square) or absence (white square) of CT, TCP, O1/O139 antigen, and the Aux3^E^ or Aux3^P^ module is indicated. Environmental (yellow), O1 Classical (green), Pre-7th Pandemic O1 El Tor (light blue), 7th Pandemic O1 El Tor (dark blue), and O139 (red) strains are highlighted.

### Both Aux3^P^ and Aux3^E^ Excise to Form a Circular DNA Element by Site-specific Recombination

A BLASTP search for the Aux3 integrase amino acid sequence returned a conserved domain hit for “integrase P4”, a common integrase in temperate phages and pathogenicity islands known to catalyze integration and excision (30, 31, 38, 39). During excision, recombination occurs between *attL* and *attR* to reform *attC* at the chromosomal excision junction and *attP* on the excised circular DNA element (Fig. 4*A*) (40, 41). Thus, we aimed to determine if Aux3 excises from the genome to form a circular element. We tested this hypothesis by inverted PCR with primers outside of the *att* sites (P1/P4) and primers inside the *att* sites facing outward (P2/P3 or P2.2/P3.2) (Fig. 4*A*). With this design, P1/P4 will only be brought into proximity for amplification upon excision and P2/P3 will only be in the right orientation upon circularization. Two Aux3^E^ strains (AM-19226 and 1154-74), three Aux3^P^ strains (N16961, C6706, and A1552), and two Aux3-naïve strains (DL4215 and DL4211) were tested for excision/circularization. After 4 hours of growth, all Aux3 strains excise this element from the genome (Fig. 4*B*). A band indicative of excision is also evident in the tested environmental strains due to the identical nature of the Aux3-naïve and Aux3-excised states. Further, the circular Aux3 module was present in all Aux3-encoding strains and absent from Aux3-naïve strains (Fig. 4*B*). PCR products were validated by Sanger sequencing against the expected chromosomal and plasmid excision junctions (*SI Appendix*, Fig. S6*A*).

**Figure 4.**
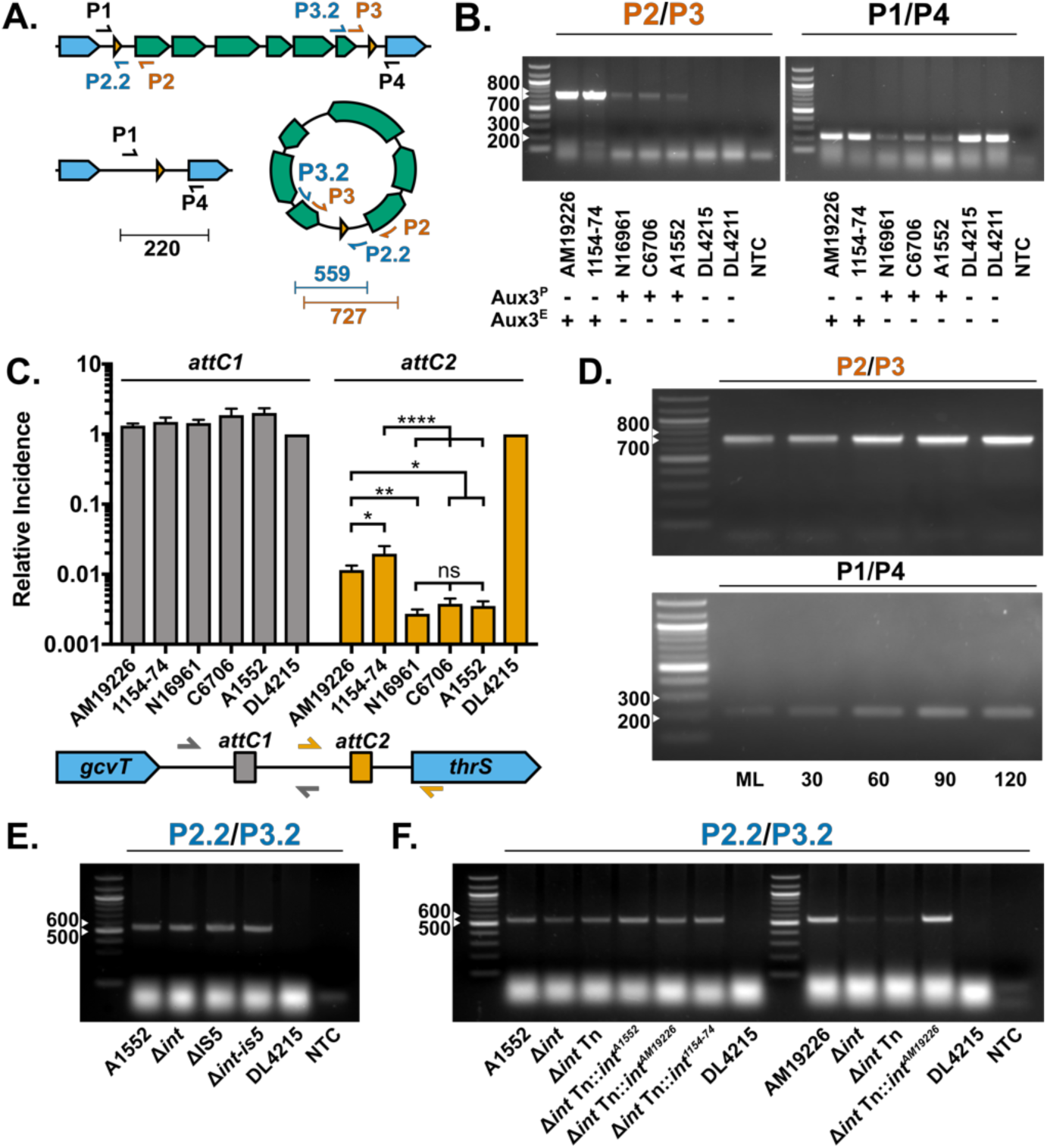
Both Aux3^E^ and Aux3^P^ modules excise from the chromosome to form a circular intermediate. (*A*) Inverse PCR schematic showing integrated and excised Aux3^P^. Aux3 genes are green, genomic flanks are blue, and *att* sites are orange triangles. Primers are represented by arrows and expected band sizes are shown. (*B*) PCR amplification of excision junctions, *attP* (P2/P3) and *attC* (P1/P4), on Aux3^E^ (AM-19226, 1154-74), Aux3^P^ (N16961, C6706, A1552), and Aux3-naïve (DL4215, DL4211) strains. (*C*) Quantification of Aux3 excision by qPCR with primers designed against the naïve *att* sites *attC1* and *attC2* (*SI Appendix*, Fig. S6*B*) on gDNA from Aux3^E^ and Aux3^P^ strains. Incidence of excision levels were determined by absolute quantification and normalization to DL4215. Error bars indicate ± SD. Significance was determined by a one-way ANOVA with Tukey’s multiple comparisons test (ns = non-significant, * = p < 0.05, ** = p < 0.01, **** = p < 0.0001). (*D*) PCR amplification of excision junctions, *attP* (P2/P3) and *attC* (P1/P4) on AM-19226 (Aux3^E^) DNA over a time course from mid log (ML) to ML + 120 min. Samples were normalized to 10ng at each time point. (*E*) Circular (P2.2/P3.2) excision junction PCR from crude DNA extracts of A1552 (Aux3^P^) wildtype, A1552 single and double recombinase null mutants, and DL4215. (*F*) Circular (P2.2/P3.2) excision junction PCR from crude DNA extracts of A1552 (Aux3^P^) and AM-19226 (Aux3^E^) wildtype strains, associated *int*-null mutants, and DL4215. Null mutants from each strain were trans-complemented with an empty mTn7 (Tn), mTn7 with the native integrase, or mTn7 with the opposing Aux3-type integrase (“integrase swap”). For all gel images, white arrows indicate ladder band sizes.

To assess the likelihood of Aux3 module transfer to a naïve strain, we measured the incidence of Aux3 excision in each strain by quantitative PCR (qPCR). Primers were designed against the Aux3-naïve *attC* sites to amplify either *attC1* or *attC2*. This experimental setup allows us to quantify excision (reversion to the naïve state) at each site (*SI Appendix*, Fig. S6*B*). With two *att* sites in the intergenic flanks, there are two potential integration states of Aux3. Measuring the reversion to a naïve site at both *attC1* and *attC2* allows us to determine which *attC* site is dynamic. Our results show that when normalized to incidence in DL4215 (Aux3-naïve or 100% excised), *attC1* is present at a ratio of approximately 1 in all tested strains (Fig. 4*C*), indicating that *attC1* is constant. The incidence of *attC2* when normalized to DL4215 is ∼1/100 genomes for Aux3^E^ strains and ∼1/500 genomes for Aux3^P^ strains (Fig. 4*C*), supporting *attC2* as the site of recombination. It is important to note that primer design was restricted by the proximity of the *attC* sites (Fig. 1*B*), leading to an optimal primer efficiency of ∼75%. While this is below the desired efficiency, it suggests that the quantification of excision is an underestimation of the true incidence. Time course analysis was performed to assess changes in excision during progression to stationary phase. Strain AM-19226 (Aux3^E^) was sampled at mid log (ML), ML+30min, ML+60min, ML+90min, and ML+120min. Quantity of DNA was normalized across time points and taken for PCR. Band intensity increases over the time course for both excision and circularization (Fig. 4*D*), indicating that excision increases with progression into stationary phase.

### Aux3^E^ and Aux3^P^ Strains Differentially Catalyze Excision and Circularization of Aux3

To investigate the role of the Aux3^P^-encoded *int* and *is5* recombinases in modular excision, each recombinase was deleted from the A1552 chromosome. Aux3^P^ circularization was assessed by inverted PCR with primers over the circular junction. New circularization primers (P2.2/P3.3) were designed because the original P2 primer binds within the deleted integrase sequence (Fig. 4*A*). Interestingly, neither single recombinase deletion nor a double knockout abrogated circularization of the Aux3^P^ module in A1552 (Fig. 4*E*). This could indicate the involvement of an unidentified Aux3-extrinsic recombinase. Conversely, the deletion of the corresponding *int* gene in the Aux3^E^ strain AM-19226 largely suppressed modular circularization, and trans-complementation of the Aux3^E^ int gene restored circularization to wildtype levels (Fig. 4*F*).

These data, along with the excision qPCR (Fig. 4*C*), suggest that there are disparities in the mechanism of site-specific recombination between Aux3^P^ and Aux3^E^ strains. One potential explanation for this difference is the presence of the IS5 module in Aux3^P^. A BLASTP search for the *int* amino acid sequence predicts this protein as a P4-like integrase and tyrosine recombinase. Pairwise alignment of the amino acid sequences of pandemic and environmental *int* proteins with other known tyrosine recombinases shows that both have all appropriate catalytic residues intact and strong homology to each other (*SI Appendix*, Fig. S7*A*). At the C-terminus, however, the Aux3^E^ integrase protein is significantly longer than the Aux3^P^ homolog. Closer investigation revealed that the IS5 element in Aux3^P^ inserted immediately downstream of the catalytic Y375 residue, blunting the true C-terminal tail of the protein and adding seven nonsense residues encoded by the 5’ end of the IS5 element (*SI Appendix*, Fig. S7*B*). We generated a predictive model of both the full-length and truncated integrase (*SI Appendix*, Fig. S7 *C* and *D*). While the orientation of the catalytic residues is unaffected, IS5 blunting results in a short, disordered C-terminal tail compared to two tyrosine-rich α-helices in the full-length protein (*SI Appendix*, Fig. S7 *C* and *D*). This could explain the decreased incidence of excision seen in Aux3^P^ strains. Counter to this idea, trans-complementation of the Aux3^E^ integrase into A1552 Δ*int*::Kan does not appear to raise Aux3^P^ excision to environmental levels (Fig. 4*F*). This suggests that the incidence of excision is not solely reliant on integrase structure, but might be differentially regulated in pandemic strains.

### Circular Aux3 Integrates Into an Aux3-naïve Chromosome at *attC2* in an *int-*dependent Manner

To assess the ability of Aux3 to integrate into the chromosome of an Aux3-naïve *V. cholerae* strain, we performed conjugative transfer experiments with an Aux3-null *V. cholerae* recipient strain and a donor *E. coli* S17 λpir carrying a pKNOCK suicide vector with variable *attP* sites (*SI Appendix*, Fig. S8 *A* and *B*). To generate a recipient strain, we first replaced the Aux3 locus in *V. cholerae* A1552 with a naïve *attC* site from environmental strain DL4211 (A1552 ΔAux3). Next, we introduced a FLAG-tagged copy of either the Aux3^P^ or Aux3^E^ integrase back into the chromosome under the control of the P_BAD_ promoter on the mini Tn7 transposon (Tn::*int*^*P*^ or Tn::*int*^*E*^), allowing us to induce integrase expression with the addition of arabinose to the culture media. Integrase expression was confirmed in these strains by western blot (*SI Appendix*, Fig. S8 *C* and *D*). It is important to note that the Aux3^P^ integrase construct is expressed at much lower levels than the Aux3^E^ integrase despite robust expression from the parental plasmid in *E. coli* (*SI Appendix*, Fig. S8*D*). The P_BAD_ promoter region and FLAG-*int*^*P*^ construct were verified by Sanger sequencing, and the reduced expression phenomenon was observed in 7 individual clones (data not shown). It is possible that the truncated *int*^*P*^ is being targeted for degradation. Aux3 donor constructs were generated in pKNOCK-Kan to either carry a stretch of circular Aux3 with both *attP1* and *attP2* intact (pKNOCK-*attP*^WT^) or a deletion of the *attP2* site (pKNOCK-*attP*^KO^) (*SI Appendix*, Fig. S8*B*). This experimental design allowed us to determine which integrase can catalyze integration of the Aux3 element into the naïve chromosome and if the recombination happens in a site-specific manner.

After 24hr co-culture of donor and recipient under inducing or repressing conditions, mixtures were plated on LB agar with kanamycin (donors), rifampicin/gentamycin (recipients), and rifampicin/gentamycin/kanamycin (transconjugants) to determine respective colony forming units (CFUs) and conjugation frequencies. For all conditions, no significant difference was seen in the recipient or donor counts (Fig. 5*A* and *SI Appendix*, Table S5). While all other conditions resulted in conjugation frequencies at or slightly above the limit of detection, transfer of pKNOCK-*attP*^WT^ to the induced Tn::*int*^*E*^ recipient resulted in a 3-log increase of conjugation frequency over baseline (Fig. 5 *B* and *C*, and *SI Appendix*, Fig. S8*E* and Table S5). The locus of integration was confirmed by PCR with P1/P4 (Fig. 5*D*), as an integrated pKNOCK-*attP*^WT^ results in a 3 kb fragment compared to the 220 bp Aux3-naïve fragment. These results demonstrate that the Aux3^E^ integrase is capable of catalyzing recombination between *attP2* on circular Aux3 and *attC2* on the naïve chromosome, and further indicate that Aux3^E^ is a MGE circulating in the aquatic *V. cholerae* reservoir.

**Figure 5.**
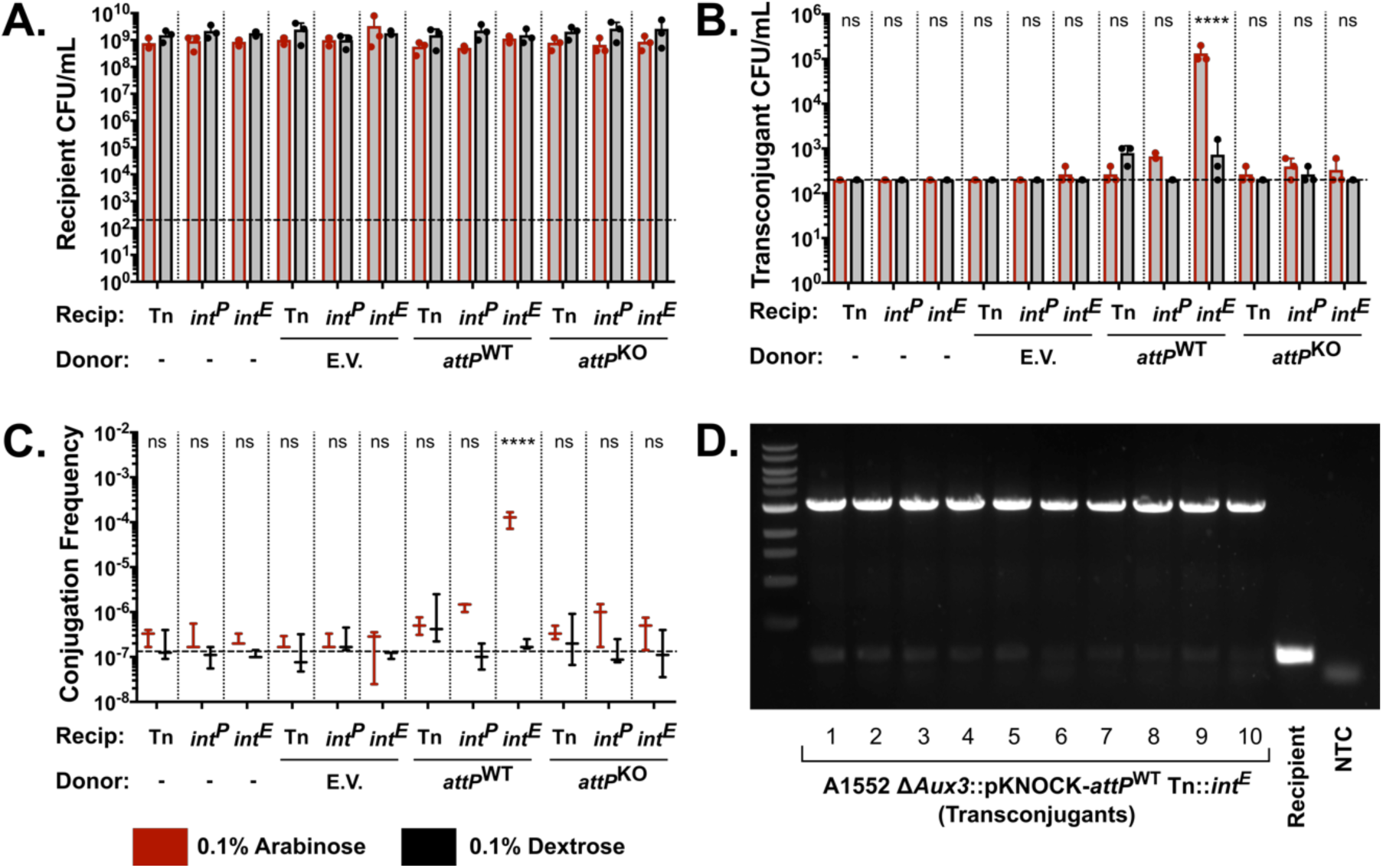
The Aux3^E^ integrase catalyzes integration of circular Aux3 into the naïve *attC* site. (*A*) Quantification of viable counts of total recipient *V. cholerae* cells (Rif^R^/Gent^R^) from conjugative transfer experiments. (*B*) Quantification of viable counts of transconjugant *V. cholerae* cells (Rif^R^/Gent^R^/Kan^R^) from conjugative transfer experiments. (*C*) Conjugation frequency from transfer experiments as determined by transconjugant counts divided by total recipient counts. (*D*) PCR verification of pKNOCK-*attP*^WT^ integration at the defined Aux3 locus with primers P1 and P4 (Fig. 4*A*). For all quantitative experiments arabinose-induced experiments are shown in red and dextrose control experiments are shown in black. Horizontal dashed line indicates the limit of detection. All quantitative experiments were performed in triplicate. Horizontal bars represent the mean and error bars indicate ± SD. Significance was determined by a 2 way ANOVA with Sidak’s multiple comparisons test (ns = non-significant, **** = p < 0.0001). E.V. = S17 λpir;pKNOCK-Kan; *attP*^WT^ = S17 λpir;pKNOCK-*attP*^WT^; *attP*^KO^ = S17 λpir;pKNOCK-*attP*^KO^; Tn = A1552 ΔAux3 Tn; *int*^*P*^ = A1552 ΔAux3 Tn::*int*^*P*^; *int*^*E*^ = A1552 ΔAux3 Tn::*int*^*E*^.

## Discussion

Here, we demonstrated that the T6SS Aux3 module is specific mainly to pandemic strains of *V. cholerae*. We further revealed that this locus is the evolutionary remnant of a prophage-like element circulating in the environmental reservoir of non-pathogenic *V. cholerae* strains. The Aux3^E^ element uses its encoded phage integrase to catalyze site-specific recombination at the flanking *att2* sites, forming a circular Aux3 element that is likely primed for horizontal gene transfer to an Aux3-naïve strain of *V. cholerae*. We showed that this locus is partially conserved and expanded within the pandemic lineage of *V. cholerae*. Despite the lack of the majority of its prophage structural and regulatory genes in the pandemic Aux3^P^, this locus maintains its P4-like integrase and flanking *att* sites. Site-specific recombination of this locus is conserved at lower levels in pandemic strains, although the Aux3 integrase does not seem to be necessary for this process. Finally, we show that the Aux3^E^ integrase is capable of integrating a circular Aux3 element into an Aux3-naïve chromosome at the *attC2* site.

The T6SS is a vital defense mechanism for *V. cholerae* and several other pathogenic Gram-negative species in both the host pathogenesis process and interbacterial competition. It is hypothesized that the T6SS is an evolutionary repurposing of a bacteriophage infection (26, 27), but the system is conserved so far back in the *Vibrio* lineage that we have not seen evidence of the initial prophage infections that evolved into the system as it exists today. We believe that our findings offer a snapshot of early T6SS evolution, in which a lysogenic phage infection was degraded to solely the components necessary to increase host fitness. Our results indicate that the pandemic Aux3 locus in *V. cholerae* is related to an environmentally circulating phage-like element that possibly degraded to form the six-gene pandemic-specific module. The route of transfer in the environmental reservoir is currently unknown, as we have no experimental data to support Aux3^E^ producing its own phage particle. Two potential mechanisms by which Aux3^E^ could be transferred between strains without making its own phage particle are generalized transduction by environmental lytic phages or chitin-induced natural competence. For the latter mechanism, excision and circularization would likely confer no advantage, as the *V. cholerae* natural competence machinery can foster the transfer of linear genome fragments significantly larger than the Aux3^E^ module (42). Several *V. cholerae* modules capable of site specific-recombination, however, are transferred by lytic phage transduction (43–45), and it is likely that the circular intermediate is more readily packaged into the transducing phage particle.

In *V. cholerae*, a major role for the T6SS is intraspecies competition and intra-host survival, and the acquisition of new effector proteins could be a key factor in a strains’ success or failure in these processes. The phenomenon of T6 effector exchange in *V. cholerae* has been highlighted (19, 20, 24), but the mechanism has remained elusive. Here we describe a site-specific recombination mechanism of T6SS effector acquisition for the Aux3 locus. The acquisition of genomic islands by this mechanism is not uncommon in *V. cholerae* (29–32). For instance, the GI*Vch*S12 element encodes its own integrase, excises from the chromosome to form a circular element, and carries a cluster of T6SS genes including an *hcp* gene and an E/I pair (32). Aux3, however, can be differentiated from GI*Vch*S12 by its distribution. Like Aux3^E^, GI*Vch*S12 circulates in the environmental reservoir of *V. cholerae* by apparent HGT (20), but Aux3 expands into the pandemic lineage. This indicates that the acquisition of Aux3^E^ and the eventual reduction to Aux3^P^ may have been an important step in the transition from environmental to pandemic organism.

Further supporting the potential fitness advantage of Aux3, our results show a disparity in the quantity of excision between Aux3^E^ and Aux3^P^. Truncation of *int*^*E*^ appears to have occurred by insertional sequence (IS5 element) interruption to form *int*^*P*^. IS5 elements have been shown to drive rapid adaptation in response to environmental stress through either transcriptional regulation of nearby genes or through insertional inactivation (46–49). Here we show that the IS5-truncated *int*^*P*^ is expressed at much lower levels than the full-length *int*^*E*^, despite having the same promoter and induction conditions. We speculate that truncation of the *int* gene by IS5 leads to degradation of the Int protein and reduced excision or integration of Aux3. Whether this degradation occurs non-specifically due to truncation and improper folding or as a specific consequence of the short C-terminal tail added by the IS5 element remains to be shown. Excision also appears to be differentially regulated in pandemic strains. Transcomplementation of *int*^*E*^ into pandemic *V. cholerae* did not increase excision to Aux3^E^ levels. Over-expression of *int*^*E*^ in pandemic *V. cholerae* did, however, catalyze increased integration of our Aux3 surrogate vector, while *int*^*P*^ over-expression did not. Regulation of Aux3 recombination in pandemic strains may have been re-wired to favor the integration of Aux3 over excision. Genes encoded by the Aux3 element may have conferred a competitive edge to a common ancestor of the pandemic clade, and this advantage was locked into the chromosome by IS5 insertion and shifts in overall regulation of excision. This biological phenomenon is referred to as “phage grounding”, and is highly advantageous to the host cell harboring the newly inactivated lysogen (50). This is, in fact, one potential route by which the T6SS itself was first acquired. In the case of Aux3, this advantage would likely be conferred by encoding an extra T6SS effector set, but the role of *tseH* in T6SS-dependent competition is still unclear (33). We believe that our findings yield several indications that Aux3 integration was selected for during the evolution of pandemic *V. cholerae* and that further mechanistic studies of the role of *tseH* are warranted.

## Materials and Methods

*SI Appendix, Supplementary Materials and Methods* describes in detail the procedures used in this study, including the following: bacterial growth conditions, mutant strain construction, identification of bacteriophage elements, sequence alignment, Aux3 enrichment analysis, phylogeny and tree building, excision PCR and qPCR, and Aux3 module transfer experiments.

## Acknowledgments

We acknowledge Michelle Dziejman, David Rozak, and Melanie Blokesch for strains and plasmid constructs necessary for the completion of this study. This work was supported by the National Institutes of Health (1R01AI139103-01A1) and the University of Colorado Anschutz Medical Campus.

