## Supplemental Data for "*Vibrio cholerae* Type VI Secretion System Auxiliary Cluster 3 is a Pandemic-associated Mobile Genetic Element"

* Stefan Pukatzki

### Supplementary Materials and Methods

#### Bacterial Strains, Plasmids, and Growth Conditions

*V. cholerae* strains, *E. coli* strains, and plasmids used in this study are listed in Table S1. *E. coli* strains DH5α λpir (1), SM10 λpir (2), and S17 λpir (2) were used for cloning and as donor strains in conjugative transfer experiments. All strains were routinely cultured at 37 ºC in Lysogeny Broth (LB) with shaking at 250 rpm. Culture on agar plates was done on LB agar at 37 ºC or 30 ºC. When required, arabinose/dextrose (for the expression or repression, respectively, of *int* under the control of the $P_{BAD}$ promoter) or antibiotics were added to both liquid or agar culture medium at the following concentrations: 0.1% arabinose, 0.1% dextrose, 100 µg/mL ampicillin, 100 µg/mL streptomycin, 100 µg/mL spectinomycin 50 µg/mL rifampicin, 50 µg/mL kanamycin, and 10 µg/mL gentamycin. Instant Ocean (7g/L IO) and chitin flakes (8 g/150 mL, MP Biomedicals) were used for MuGENT cloning experiments.

#### Bacterial Mutant Strain Construction, Plasmid Construction, and Trans-complementation

All DNA manipulations were performed according to standard molecular biology protocols. The following enzymes/kits were used according to manufacturer specifications: Phusion High-Fidelity DNA Polymerase (Thermo Fisher), restriction enzymes (New England Biolabs), NEBuilder HiFi DNA Assembly Cloning Kit (New England Biolabs), and Taq PCR Master Mix (Qiagen). Engineered plasmids and bacterial strains were verified by colony PCR and Sanger sequencing (Quintara Biosciences).

Genetic deletion in *V. cholerae* were generated by either allelic exchange and sucrose counter selection with the suicide vector pCVD442 (3) or the natural transformation-based MuGENT method (4). The plasmid pKD13 (5) served as template for the amplification of the flippable antibiotic cassette FRT-Kan-FRT. For allelic exchange, the flippable Kan cassette was inserted between 1 kb homology arms in pCVD442 by Gibson cloning (6), and clones were selected by sucrose counterselection and kanamycin resistance. The MuGENT technique was used to generate A1552 ΔIS5 and A1552 ΔAux3. Unmarked ΔIS5 construct was generated by Gibson cloning of 3kb fragments upstream and downstream of the Aux3 IS5 element into pUC19 and amplification of a 6kb fragment with pUC19 specific primers. Unmarked ΔAux3 construct was generated by amplifying a 6kb fragment containing naïve Aux3 *attC* site from the DL4211 chromosome. Selective fragments were generated by Gibson cloning a spectinomycin resistance cassette from SAD033 in between 3kb homology arms encompassing the *V. cholerae lacZ* gene and the surrounding sequence into pUC19. The 5.8kb selective *lacZ*::Spec fragment was amplified using primers ABD334/ABD335 (4). Unmarked and selective fragments were co-transformed into *V. cholerae* cells on chitin. White/spectinomycin resistant cells were screened for the unmarked mutation. The spectinomycin resistance cassette was cured from *lacZ* with pCVD442 carrying a wildtype copy of *V. cholerae lacZ*.

All transcomplementation vectors were generated by Gibson cloning. Expression constructs with either endogenous or $P_{BAD}$ promoters were inserted into the mini Tn7 transposon (mTn7/Tn) in pGP704-mTn7. Constructs were moved onto the *V. cholerae* chromosome by tri-parental mating (7–9).

Donor pKNOCK-*attP* vectors were generated by Gibson cloning. The circular Aux3 *attP* region was also generated by Gibson cloning. Primers overlapping the *att* site were modified to remove the *attP2* site. Regions containing the *attP* sites were amplified off the Gibson assembled fragments and assembled into the SmaI-cut pKNOCK-Kan vector.

**Identification of *att* Sites and Bacteriophage Elements**

To identify potential *att* sites, intergenic sequences from VCA0280 to VCA0281 and VCA0286 to VCA0287 were concatenated and input into REPFIND (10) with a minimum repeat size of 10 and a P-value cutoff of 0.0001. To identify putative prophages, GenBank files for $Aux3^{E}$ strains were submitted to PHASTER (11).

**Nucleotide/Amino Acid Sequence Alignment**

All genomes used for alignments can be found in *SI Appendix,* Table S3. All nucleotide alignments outside of phylogenetic analyses were performed in Geneious Prime (v2019.0.4). Nucleotide sequences encompassing more than one open reading frame were aligned using the Progressive MAUVE algorithm (12) to account for insertions, deletions, and rearrangements. Single gene, intergenic region, or single protein sequence pairwise alignments were performed using MUSCLE (v3.8.425) (13).

**Aux3 Enrichment Analysis**

All Megablast queries were performed in Geneious Prime (v2019.0.4). All downstream manipulations and plots were done in RStudio (R version 3.3.2 (2016-10-31) -- "Sincere Pumpkin Patch"). Genomic FASTA files for 749 *V. cholerae* strains and closely related species were downloaded from the PATRIC database (14). Nucleotide sequences for *tseH* (VCA0281), *tseL* (VC1418), *vasX* (VCA0020), *vgrG3* (VCA0123), *tcpA* (VCA0828), and *ctxAB* (VC1456-VC1457) from O1 El Tor type strain N16961 were queried by Megablast against a custom database of the downloaded PATRIC sequences to generate a grade (a weighted metric combining query coverage (.50), e-value (.25), and pairwise identity (.25)) for each gene locus in each strain. Strains were grouped based on a 99% grade cutoff for *tseH* and the three A-type effectors *tseL*, *vasX*, and *vgrG3* to create 4 groups (*SI Appendix*, Table S4) and assess co-occurrence of *tseH* in AAA pandemic strains by Fisher's Exact Test.

PATRIC strains were k-means clustered by Partitioning Around Medoids (pam, R package cluster v2.1.0) based on grades for *tseH*, *tseL*, *vasX*, *vgrG3*, *ctxAB*, and *tcpA*. Mean grade was determined at each locus for each cluster and plotted as a heat map (pheatmap, R package pheatmap v1.0.12).

**Phylogenetic Analysis and Tree Building**

Genomic FASTA files for tree building were obtained from the PATRIC database (14) or NCBI and annotated using Prokka (v1.12) (15). A core genome was extracted from Prokka-output GFF3 files and aligned using Roary (v3.11.2) (16). The core genome alignment was reduced to loci harboring polymorphisms using SNP-sites (v2.4.1) (17). Phylogenetic tree was built using the RAxML GTR Gamma Maximum Likelihood model. Statistical branch support was obtained from 100 bootstrap repeats. Phylogenetic trees were visualized from RAxML-generated newick files using TreeGraph 2 (v2.15.0-887 beta) (18). Branches with bootstrapping support values <70 were collapsed. Presence of TCP and CTX were determined by Megablast for *tcpA* (VC0828) and *ctxAB* (VC1456-VC1457). O1 antigen status was determined from the literature. Presence of *tseH* was determined as described above.

**Excision/circularization PCR and quantitative PCR**

Bacterial strains analyzed by excision/circularization PCR or qPCR were grown overnight as described above. Approximately equivalent growth for all analyzed strains was verified (*SI Appendix*, Fig. S6*C*). For Fig. 4 *B*, *E*, and *F* overnight cultures were subcultured 1:50 in 5 mL of fresh LB and grown for 4 hours at 37 ºC with shaking at 250 rpm. Cultures were pelleted (4,300 xg, 10 min) and resuspended at 10X concentration in nuclease free $H_2O$. Cell suspensions were boiled for 5 min to release nucleic acids. PCR was performed on 3 µL of each lysate with the indicated primers. 3% DMSO was added for reactions using P2.2/P3.2 due to lower primer efficiency. For Fig 4*D* overnight cultures were subcultured 1:50 in 10 mL of fresh LB and grown to mid log (OD600 = 0.4). 1 mL of culture was collected and pelleted (14,000 rpm, 2 min) at mid log and 30 min intervals post mid log. Plasmid DNA was purified with the NEB Monarch Plasmid Miniprep Kit. All samples were normalized to 10 ng/µL, and PCR was performed on 1 µL of each prep with the indicated primers. For excision/circularization qPCR, overnight cultures were subcultured 1:50 in 5 mL of fresh LB and grown for 4 hours at 37 ºC with shaking at 250 rpm. 1 mL of culture was collected and pelleted (14,000 rpm, 2 min), and DNA was extracted by phenol/chloroform extraction. DNA for all samples was normalized to 20 ng/ µL. qPCR was performed on 5 µL (100 ng) of each sample in a 20 µL reaction volume with Bio-Rad SYBR Green Master Mix according to the product manual. PCR was performed with primers against *attC1*, *attC2*, and *ompW*. All targets were measured by absolute quantification against a standard curve of DL4215 genomic DNA (Aux3-naïve). *attC* site quantity was normalized to *ompW* to control for variability in the quantity of input DNA. Finally, all *ompW*-normalized samples were normalized to DL4215 samples to determine the ratio of excised genomes as DL4215 is in a "permanently Aux3-excised" state. Averages of at least three independent experiments (± standard deviation) are provided.

**Aux3 Module Transfer Experiments**

Overnight cultures of recipient strains (A1552 ΔAux3 with variable mTn7 constructs) and donor strains (S17 λpir with variable pKNOCK vectors) were pelleted (4,300 xg, 10 min) and resuspended at 10X concentration in LB media. 10 µL of each concentrated cell suspension was resuspended in 1 mL LB (1:100), from which 7 x 1:10 serial dilutions were prepared. Serial dilutions were plated as 5 µL spots on LB agar to determine input colony forming units. Donor and recipient strains were mixed in all combinations at a ratio of 10:1 donor to recipient. Mixtures were plated at 25 µL spots on Durapore .22 µm PVDF filters (Millipore Sigma) on pre-dried, pre-warmed LB agar plates with either arabinose or dextrose. Spots were dried and incubated at 37 ºC for 24 hours. After 24 hour incubation, filters were collected and submerged in 1 mL LB media. Bacterial cells were resuspended by vortexing. Serial dilutions (7 x 1:10) were prepared from each suspension, and 5 µL spots of each dilution were plated on LB agar with kanamycin (donors), rifampicin/gentamycin (recipients), or rifampicin/gentamycin/kanamycin (transconjugants) to determine colony forming units (CFU) for each subset of cells. CFU/mL was determined by counting colonies from the lowest countable dilution and adjusting for volume plated and dilution factor. Conjugative frequency was determined by dividing transconjugant CFU/mL by total recipient CFU/mL. Averages of at least three independent experiments (± standard deviation) are provided (*SI Appendix*, Table S5).

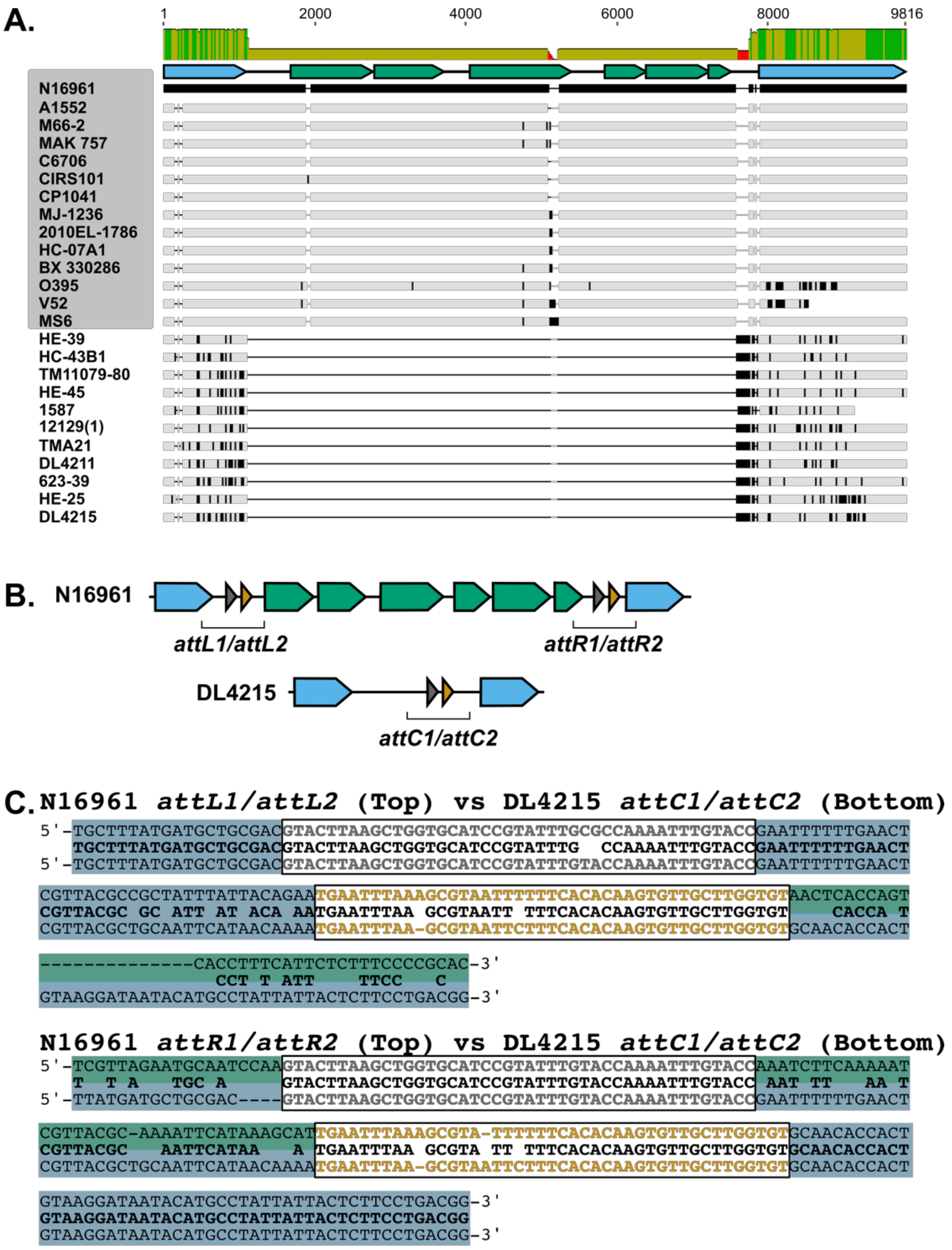


**Fig. S1.** Aux3 was likely integrated site-specifically at *attC2* in the *V. cholerae* chromosome. (*A*) MAUVE alignment of the Aux3 locus in pandemic (grey background) and environmental (white background) *V. cholerae* strains. N16961 is set as the reference sequence. Black indicates disagreement to the reference. Identity is represented in bars above the alignment (green = 100%, yellow = 99%-30%, red = <30%). Aux3 schematic is shown under the identity graph (blue = genomic flanks, green = Aux3 genes). (*B*) Schematic of Aux3 *att* sites in both the integrated (N16961) and naïve (DL4215) state. Genomic flanking genes are shown in blue. Aux3 genes are shown in green. Attachment site *att1* is shown in grey and attachment site *att2* is shown in orange. (*C*) MUSCLE alignment of the Aux3 flanking *attL* sites (top) and *attR* sites (bottom) in N16961 to the *attC* sites in DL4215. Matching bases are shown in bold black. Phage-like *att* sites are boxed and bolded (*attL1/attR1/attC1* in grey, *attL2/attR2/attC2* in orange). Genomic flanks are highlighted in blue. Aux3 module is highlighted in green.

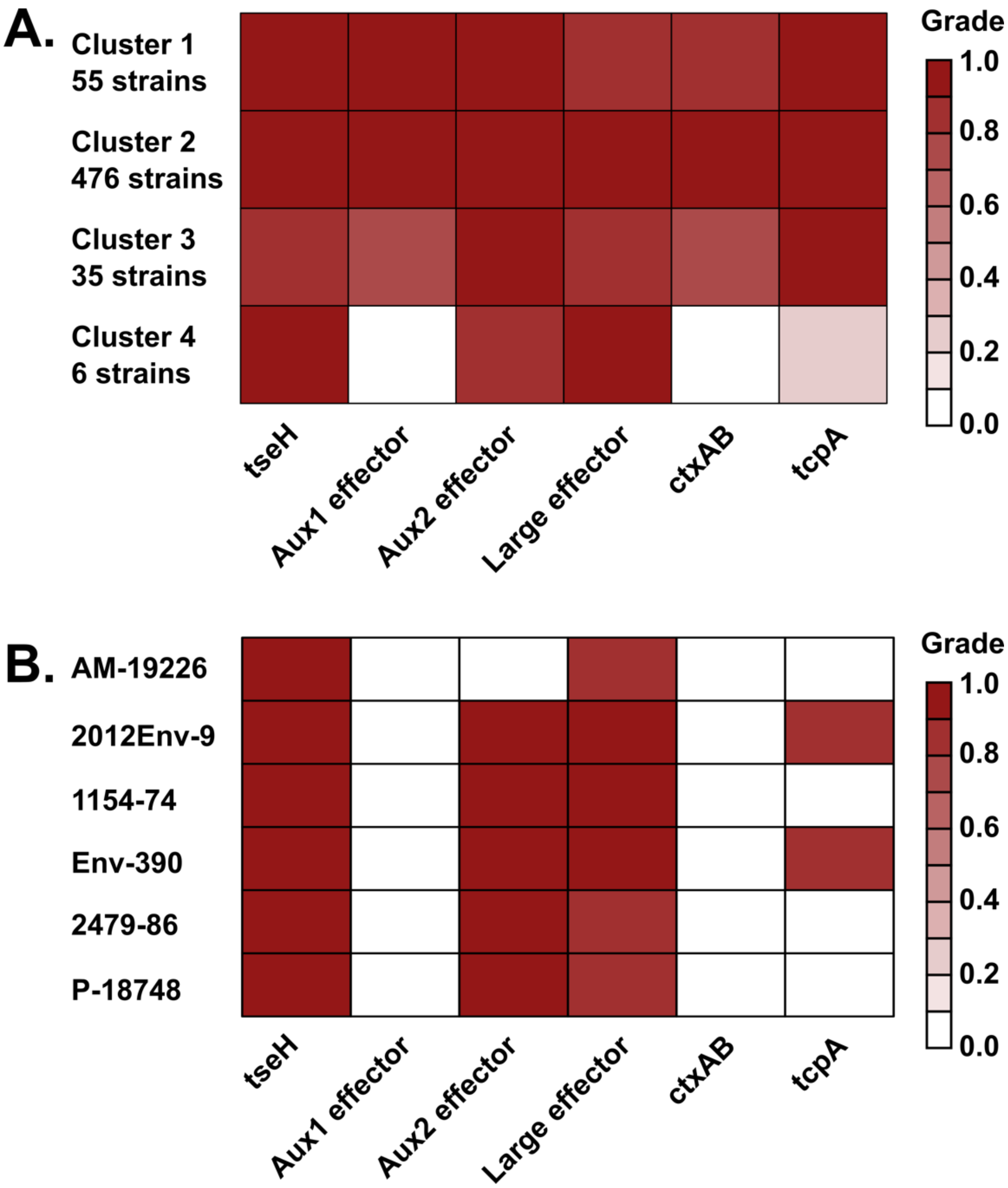


**Fig. S2.** The *tseH* gene is found in pandemic *V. cholerae* and a small environmental reservoir. (*A*) Heatmap of 572 *tseH* (+) *V. cholerae* genomes from the PATRIC database. Genomes are collapsed into clusters based on mean BLAST Grade for *tseH* and five other pandemic associated factors: *tseL* (Aux1), *vasX* (Aux2), *vgrG3* (Large), *ctxAB* (CT), and *tcpA* (TCP). (*B*) Expanded heatmap of the six strains assigned to cluster 4. Lack of *tseL*, *ctxAB*, and *tcpA* indicates that these are non-pathogenic environmental strains.

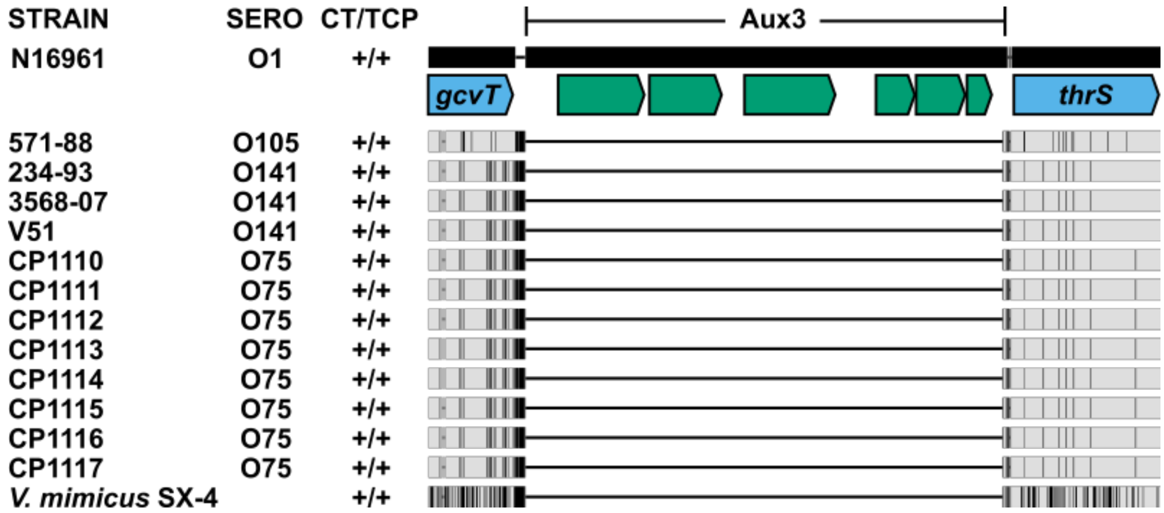


**Fig. S3.** Aux3 is absent from non-pandemic, pathogenic *V. cholerae* and *V. mimicus* strains. MAUVE alignment of the Aux3 locus (VCA0280-VCA0287) in N16961 and 13 nonO1/O139, CTX+/TCP+ *V. cholerae* and *V. mimicus* strains that have been shown to cause isolated disease. N16961 is set as a reference sequence and difference against the reference are shown below. Grey bars indicate agreements and black bars indicate differences.

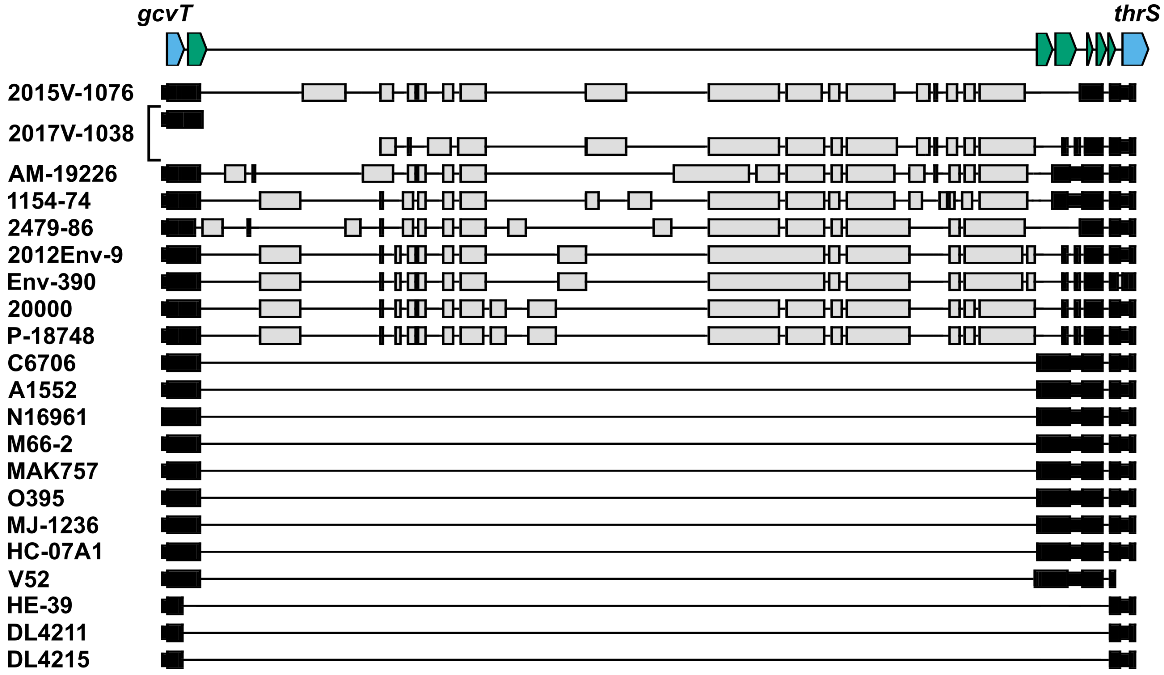


**Fig. S4.** Environmental Aux3 strains encode variable extra sequence between *int* and VCA0283. MAUVE alignment of the Aux3 locus (VCA0280-VCA0287) for 9 Aux3-encoding environmental strains, 9 Aux3-encoding pandemic strains, and 3 Aux3-naïve strains. Black bars indicate nucleotides conserved in the pandemic Aux3-encoding strains. Grey bars indicate nucleotides absent from the pandemic Aux3 element. Pandemic Aux3 schematic is shown (top) with genomic flanking genes in blue and Aux3 genes shown in green.

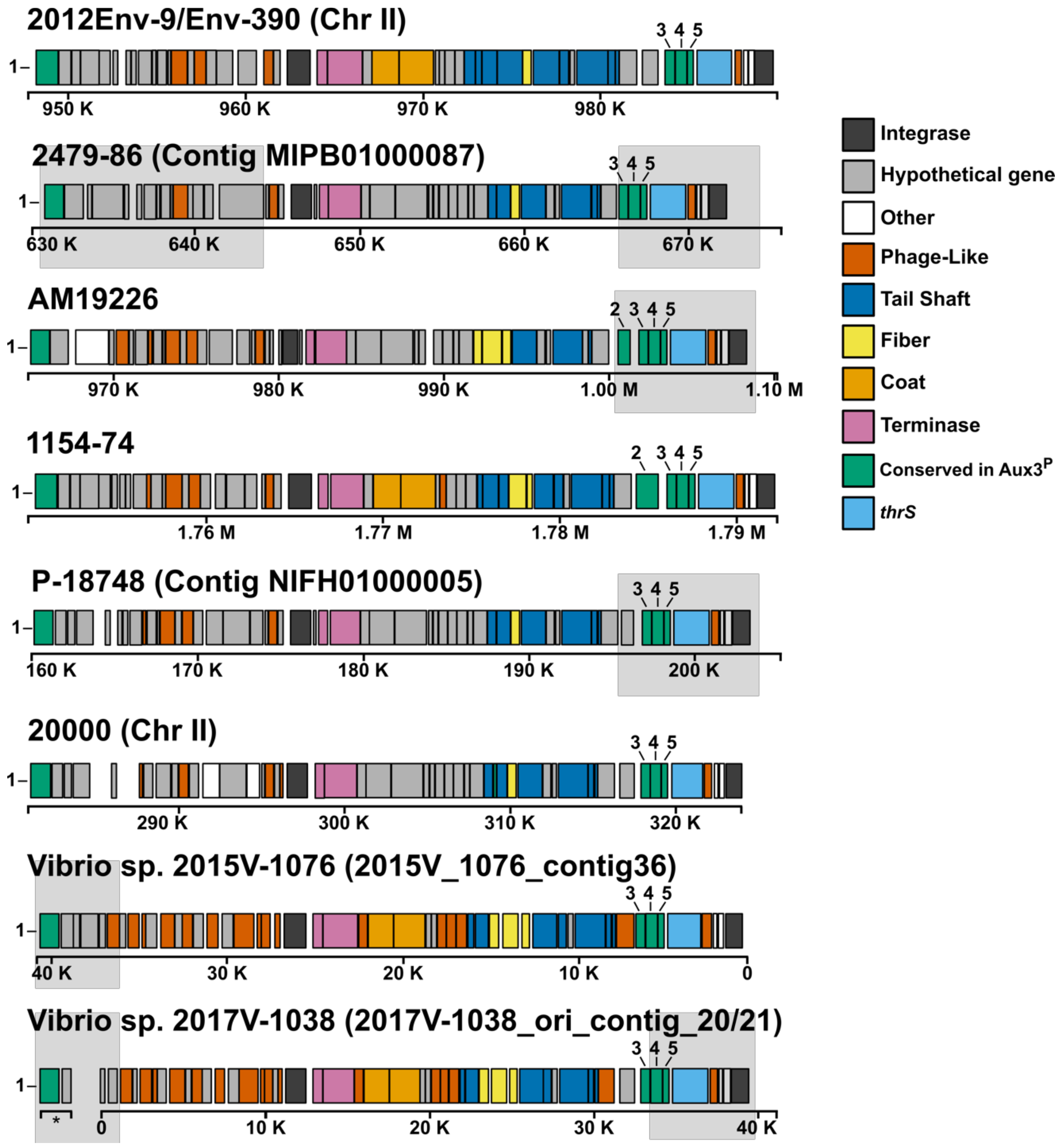


**Fig. S5.** Aux3$^E$ strains encode putatively intact prophage elements. PHASTER genome diagrams showing predicted prophage regions from *int* (VCA0281) through the Aux3-extrinsic superintegron integrase *intI4* (VCA0291). Coding regions are colored according to homology to broad categories of known phage genes. Genes conserved in the pandemic Aux3 module are indicated (1 = *int*, 2 = VCA0283, 3 = *PAAR*, 4 = *tseH*, 5 = *tsiH*). Light grey boxes indicate regions not called by PHASTER, but confirmed by sequence analysis. * indicates coding regions found on a separate contig. Strains are shown in order from most closely related to the pandemic clade to least closely related (Fig. 3).

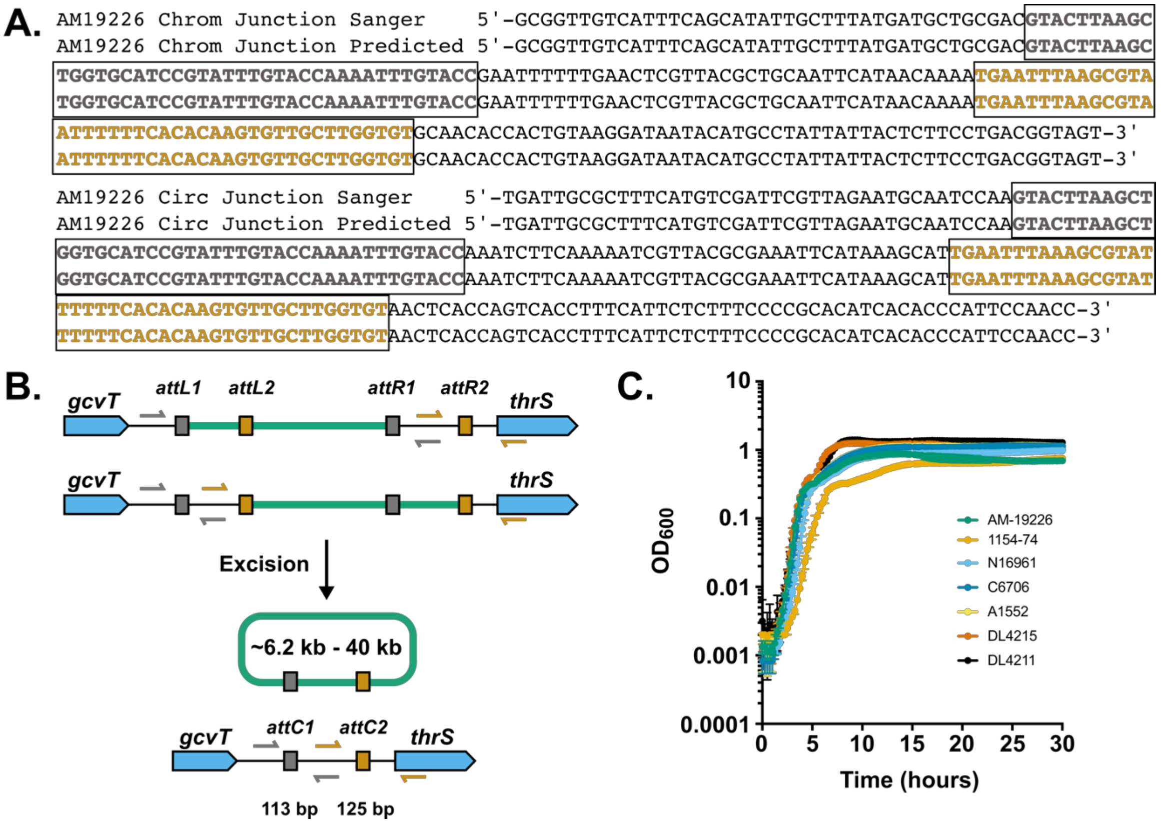


**Fig. S6.** Excision PCR and qPCR controls and schematics. (*A*) MUSCLE alignment of predicted chromosomal and circular junction post Aux3 excision with Sanger sequence of inverted PCR bands from AM-19226. (*B*) Excision qPCR schematic. Grey arrows represent *attC1* primers. Orange arrows represent *attC2* primers. Primer binding sites for the two possible states for Aux3 integration (Top) and the excised state (Bottom) are shown. (*C*) Growth curves performed in triplicate for all strains used in PCR/qPCR experiments show that all strains grow approximately the same under the experimental conditions.

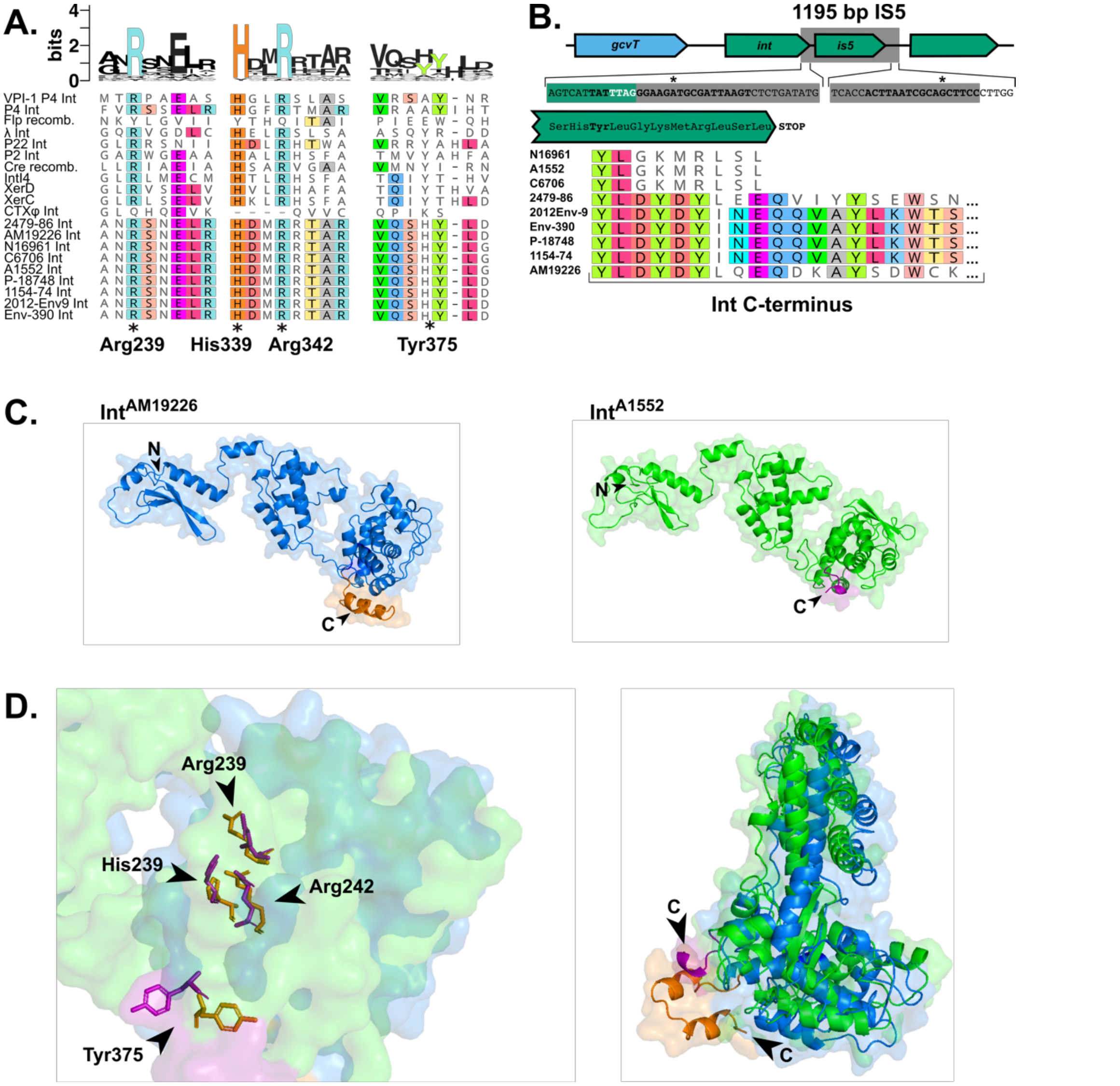


**Fig. S7.** Reduced excision in $\text{Aux3}^{P}$ may be due to integrase truncation. (*A*) MUSCLE alignment of $\text{Aux3}^{P}$ and $\text{Aux3}^{E}$ integrases with known tyrosine recombinases. Three regions containing the four main catalytic residues (19) are shown with the known catalytic residues highlighted in the SeqLogo (top), by *, and by named below. (*B*) Schematic of IS5 element insertion into and blunting of the Aux3 integrase. The IS5 element is highlighted by a grey rectangle in the gene diagram (top). The black bolded nucleotide sequence indicates the canonical inverted repeats and a * marks the known disagreement in these repeats. The white bolded text indicates the IS5 target site (Pyr-TA-Pur) (20) immediately after the catalytic Tyr (Y) residue. MUSCLE alignment (bottom) of the C-terminal amino acid sequence of $\text{Aux3}^{P}$ and $\text{Aux3}^{E}$ integrases indicates the introduction of a nonsense tail and premature STOP by the 5' sequence of the IS5 in pandemic strains. (*C*) Phyre2 (21) intensive model of $\text{Int}^{AM19226}$ (left) and $\text{Int}^{A1552}$ (right) visualized with PyMol (v1.2r3pre). C-terminal tails after the catalytic tyrosine are colored orange or magenta, respectively. N- and C-termini are highlighted by black arrows. (*D*) Overlay of surface models of $\text{Int}^{AM19226}$ and $\text{Int}^{A1552}$. Catalytic residues are orange ($\text{Int}^{AM19226}$) or magenta ($\text{Int}^{A1552}$) and highlighted by black arrows (left). Disparity in the C-terminal tail (right) is shown by a ribbon model. C-termini are highlighted by black arrows.

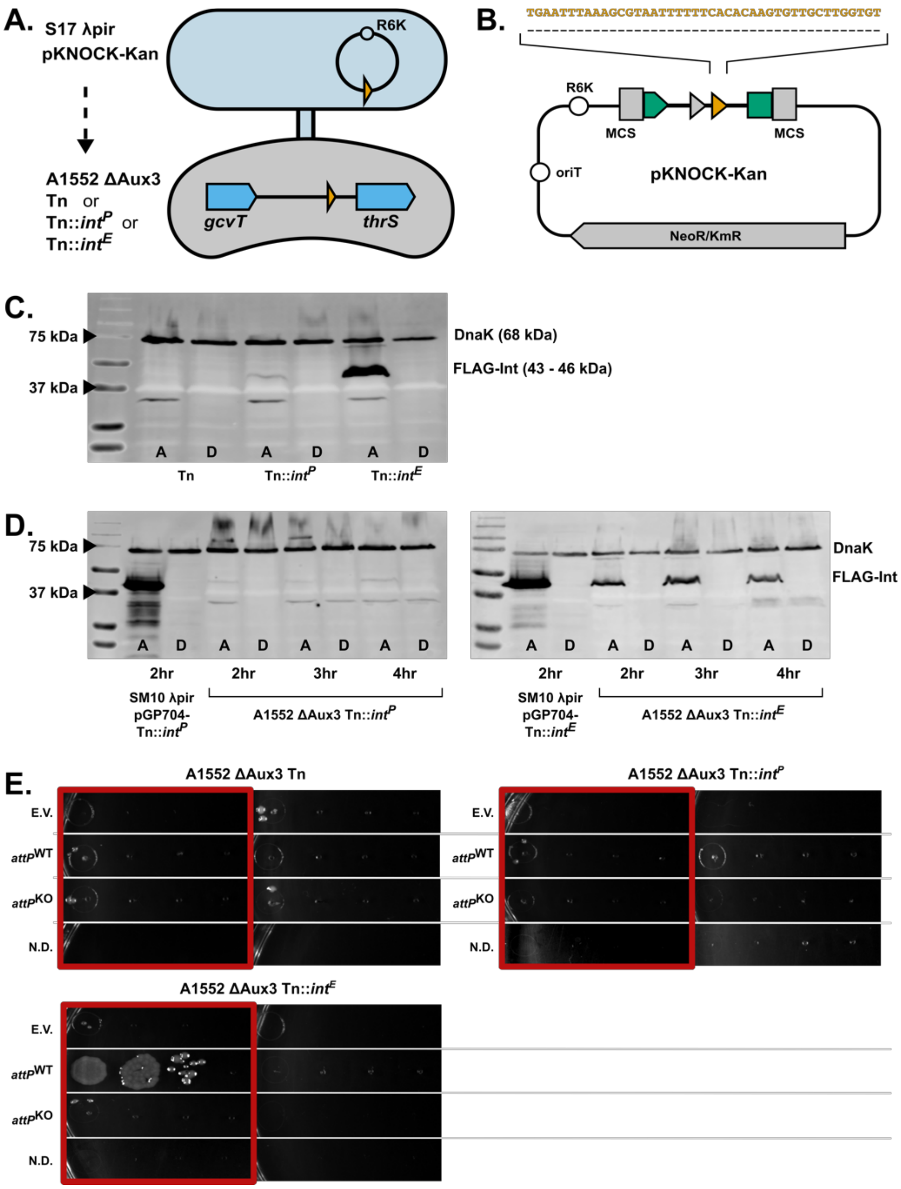


Fig. S8. Arabinose induction of *FLAG-int*$^{E}$ drives integration of pKNOCK-*attP*$^{WT}$. (*A*) Schematic of Aux3 transfer experiments showing conjugation between donor strain S17 λpir carrying variable pKNOCK-Kan vectors and recipient strain *V. cholerae* ΔAux3 with variable *int* genes expressed off of the mini Tn7 transposon. (*B*) Schematic of pKNOCK-Kan vectors carrying either WT *attP* or *attP2* knockout fragments. (*C*) Western blot analysis of 4 hr integrase induction in *V. cholerae* ΔAux3 strains. A = 0.1% arabinose. D = 0.1% dextrose. (*D*) Western blot analysis of integrase induction in *V. cholerae* ΔAux3 strains and their respective SM10 λpir parental strains over a time course from 2 to 4 hours. (*E*) Representative images from transconjugant dilution plates from Aux3 transfer experiments (Fig. 5). Images highlighted in red are from arabinose-induced experiments. E.V. = S17 λpir;pKNOCK-Kan; *attP*$^{WT}$ = S17 λpir;pKNOCK-*attP*$^{WT}$; *attP*$^{KO}$ = S17 λpir;pKNOCK-*attP*$^{KO}$; N.D. = No Donor.

**Table S1.** Bacterial Strains and Plasmids

| Strain or Plasmid | Genotype*/Description | Internal Strain Ref. | Reference |
|---|---|---|---|
| DH5α λpir | $F^-$ endA1 glnV44 thi-1 recA1 relA1 gyrA96 deoR nupG φ80lacZΔM15 Δ(lacZYA-argF) U169 hsdR17 ($r_K^-$ $m_K^+$) phoA, $λ^-$ | FJS010 | (1) |
| SM10 λpir | thi-1 thr leu tonA lacY supE recA::RP4-2-Tc::Mu, Kmr (λpir); $Kan^R$ | FJS011 | (2) |
| S17 λpir | Tpr Smr recA thi pro hsdR2M1 RP4:2-Tc:Mu:Kmr Tn7 (λpir); $Str^R$ | FJS275 | (2) |
| SAD033 | *V. cholerae* strain carrying spectinomycin resistance cassette for MuGENT cloning | FJS028 | (Ankur Dalia) |
| N16961 | WT O1 El Tor Pandemic Strain; $Aux3^P$; $Sm^R$ | FJS002 | (22) |
| C6706 | WT O1 El Tor Pandemic Strain; $Aux3^P$; $Sm^R$ | FJS005 | (23) |
| A1552 | WT O1 El Tor Pandemic Strain; $Aux3^P$; $Rif^R$ | FJS024 | (24) |
| AM-19226 Δ*endo* | O39 Clinical Isolate deleted for the type II restriction endonuclease TdeIII; $Aux3^E$; $Sm^R$ | FJS021 | (25) |
| 1154-74 | WT O49 Environmental Isolate; $Aux3^E$; $Sm^R$ | FJS038 | (26) |
| DL4215 | WT Environmental Isolate; Aux3-naïve; $Sm^R$ | FJS031 | (27) |
| DL4211 | WT Environmental Isolate; Aux3-naïve; $Rif^R$ | FJS027 | (27) |
| A1552 Δ*int*::Kan | A1552 deleted for VCA0281; $Rif^R$, $Kan^R$ | FJS189 | This study |
| A1552 ΔIS5 | A1552 deleted of the 1195 bp $IS5^{Aux3}$ element including VCA0282; $Rif^R$ | FJS138 | This study |
| A1552 Δ*int-is5*::Kan | A1552 deleted for VCA0281-VCA0282; $Rif^R$, $Kan^R$ | FJS182 | This study |
| A1552 Δ*int*::Kan Tn | A1552 Δ*int*::Kan containing empty mini-Tn7; $Rif^R$, $Kan^R$, $Gent^R$ | FJS209 | This study |
| A1552 Δ*int*::Kan Tn::$int^{A1552}$ | A1552 Δ*int*::Kan containing mini-Tn7-$P_{end}$-$int^{A1552}$; $Rif^R$, $Kan^R$, $Gent^R$ | FJS210 | This study |
| A1552 Δ*int*::Kan Tn::$int^{AM19226}$ | A1552 Δ*int*::Kan containing mini-Tn7-$P_{end}$-$int^{AM19226}$; $Rif^R$, $Kan^R$, $Gent^R$ | FJS211 | This study |
| A1552 Δ*int*::Kan Tn::$int^{1154-74}$ | A1552 Δ*int*::Kan containing mini-Tn7-$P_{end}$-$int^{1154-74}$; $Rif^R$, $Kan^R$, $Gent^R$ | FJS212 | This study |
| AM-19226 Δ*endo* Δ*int*::Kan | AM-19226 Δ*endo* deleted for VCA0281 equivalent; $Sm^R$, $Kan^R$ | FJS206 | This study |
| AM-19226 Δ*endo* Δ*int*::Kan Tn | AM-19226 Δ*int*::Kan containing empty mini-Tn7; $Sm^R$, $Kan^R$, $Gent^R$ | FJS207 | This study |
| AM-19226 Δ*endo* Δ*int*::Kan Tn::$int^{AM19226}$ | AM-19226 Δ*int*::Kan containing mini-Tn7-$P_{end}$-$int^{AM19226}$; $Sm^R$, $Kan^R$, $Gent^R$ | FJS208 | This study |
| S17 λpir; pKNOCK-$attP^{WT}$ | Conjugative *E. coli* carrying pKNOCK-$attP^{WT}$ | FJS276 | This study |
| S17 λpir; pKNOCK-$attP^{KO}$ | Conjugative *E. coli* carrying pKNOCK-$attP^{KO}$ | FJS277 | This study |
| S17 λpir; pKNOCK-Kan | Conjugative *E. coli* carrying empty pKNOCK-Kan | FJS278 | This study |

| | | | |
|---|---|---|---|
| A1552 ΔAux3 | A1552 deleted for Aux3 by homologous recombination with the DL4211 empty *attC* region (MuGENT); $Rif^R$ | FJS139 | This study |
| A1552 ΔAux3 Tn | A1552 ΔAux3 containing empty mini-Tn7; $Rif^R$, $Gent^R$ | FJS213 | This study |
| A1552 ΔAux3 Tn::$int^P$ | A1552 ΔAux3 containing mini-Tn7-*araC*-$P_{BAD}$-FLAG-$int^{A1552}$; $Rif^R$, $Gent^R$ | FJS266 | This study |
| A1552 ΔAux3 Tn::$int^E$ | A1552 ΔAux3 containing mini-Tn7-*araC*-$P_{BAD}$-FLAG-$int^{AM19226}$; $Rif^R$, $Gent^R$ | FJS267 | This study |
| **Plasmids** | | | |
| pUC19-*lacZ*::$Spec^R$ | pUC19 vector carrying spectinomycin resistance cassette inserted in between 3kb flanks homologous to *V. cholerae lacZ* and the surrounding sequence; $Amp^R$, $Spec^R$ | FJS068 | This study |
| pUC19-ΔIS5 | pUC19 vector carrying 3kb upstream and downstream flanks of the Aux3 IS5 element for IS5 deletion; $Amp^R$ | FJS085 | This study |
| pCVD442-$lacZ^{WT}$ | oriR6K, pCVD442 *sacB* counter-selectable suicide cloning vector carrying a WT copy of *V. cholerae lacZ* for curing of spectinomycin resistance cassette | FJS106 | This study |
| pKD13 | pKD vector for the amplification of FRT-flanked kanamycin cassette; $Amp^R$, $Kan^R$ | FJS159 | (5) |
| pCVD442-Δ*int*::$Kan^{FRT}$ | oriR6K, pCVD442 *sacB* counter-selectable suicide cloning vector carrying a FRT-Kan-FRT flanked by 1000 bp homology arms surrounding VCA0281 | FJS160 | This study |
| pCVD442-Δ*int-is5*::$Kan^{FRT}$ | oriR6K, pCVD442 *sacB* counter-selectable suicide cloning vector carrying a FRT-Kan-FRT flanked by 1000 bp homology arms surrounding VCA0281-VCA0282 | FJS162 | This study |
| pCVD442-Δ$int^{AM19226}$::$Kan^{FRT}$ | oriR6K, pCVD442 *sacB* counter-selectable suicide cloning vector carrying a FRT-Kan-FRT flanked by 1000 bp homology arms surrounding the AM-19226 VCA0281 homolog | FJS188 | This study |
| pUX-BF-13 | oriR6K, helper plasmid with Tn7 transposase; $Amp^R$ | FJS032 | (7) |
| pGP704-mTn7-minus SacI | pGP704 with mini-Tn7 insertion; $Amp^R$, $Gent^R$ | FJS074 | (28) |
| pGP704-mTn7-$int^{A1552}$ | pGP704 with mini-Tn7 carrying $P_{end}$-driven $int^{A1552}$; $Amp^R$, $Gent^R$ | FJS173 | This study |
| pGP704-mTn7-$int^{AM19226}$ | pGP704 with mini-Tn7 carrying $P_{end}$-driven $int^{AM19226}$; $Amp^R$, $Gent^R$ | FJS171 | This study |
| pGP704-mTn7-$int^{1154\text{-}74}$ | pGP704 with mini-Tn7 carrying $P_{end}$-driven $int^{1154\text{-}74}$; $Amp^R$, $Gent^R$ | FJS172 | This study |
| pGP704-mTn7-*araC*-$P_{BAD}$-$int^{A1552}$ | pGP704 with mini-Tn7 carrying *araC* and $P_{BAD}$-driven $int^{A1552}$; $Amp^R$, $Gent^R$ | FJS260 | This study |
| pGP704-mTn7-*araC*-$P_{BAD}$-$int^{AM19226}$ | pGP704 with mini-Tn7 carrying *araC* and $P_{BAD}$-driven $int^{AM19226}$; $Amp^R$, $Gent^R$ | FJS261 | This study |
| pKNOCK-Kan | oriR6K, knock-in plasmid; $Kan^R$ | FJS269 | (29) |

| | | | |
|---|---|---|---|
| pKNOCK-*attP*$^{WT}$ | pKNOCK-Kan with inserted 727 bp region including Aux3 *attP* amplified from pSynAux3$^{WT}$ | FJS273 | This study |
| pKNOCK-*attP*$^{KO}$ | pKNOCK-Kan with inserted 684 bp region including Aux3 *attP* amplified from pSynAux3$^{attKO}$ | FJS274 | This study |

*VC numbers as reported in (22)

**Table S2.** Primers

| Name | Sequence |
|---|---|
| P1 | GCGGTTGTCATTTCAGCATATT |
| P2 | CTGTGGCTTATCTCAGTCTTACC |
| P2.2 | GAAAGGTGACTGGTGAGTTACA |
| P3 | CAGCAACGGGTCTCAGTATT |
| P3.2 | GTGCTCTTGGCTACGTTTCT |
| P4 | TGACTACCGTCAGGAAGAGTAA |
| PattC1-F | GCGGTTGTCATTTCAGCATATT |
| PattC1-R | GAATTGCAGCGTAACGAGTTC |
| PattC2-F | CTCGTTACGCTGCAATTCATAAC |
| PattC2-R | TGACTACCGTCAGGAAGAGTAA |
| **MuGENT** | |
| *lacZ*-Upstream_pUC19-SmaI_F | tgaattcgagctcggtacccACCCTAAGCGGTTCAATTTTG |
| *lacZ*-Upstream_Spec_R | ggatccccggaatTAACGATGTGCGGGTTTTG |
| Spec_*lacZ*-Upstream_F | ccgcacatcgttaATTCCGGGGATCCGTCGAC |
| Spec_*lacZ*-Downstream_R | cggcagtgccattTGTAGGCTGGAGCTGCTTC |
| *lacZ*-Downstream_Spec_F | cagctccagcctacaAATGGCACTGCCGTACAC |
| *lacZ*-Downstream_pUC19-SmaI_R | gtcgactctagaggatccccTGATCCGATGATCTTTTCG |
| ABD334 | AGTGCTCCGGACTCTTTGCTCTG |
| ABD335 | CACTGCTCACTAGCGATGCAGTG |
| IS5-Upstream_pUC19-SmaI_F | gtcgactctagaggatccccgggGATCCTTTGGGTCACCGC |
| IS5-Upstream_R | attttccaagCTAAATAATGACTCTGAACACCAGTATG |
| IS5-Downstream_F | cattatttagCTTGGAAAATTGCGGGAG |
| IS5-Downstream_pUC19-SmaI_R | tgaattcgagctcggtacccgggAAACTGTTCAATATACGCGC |
| IS5-KO_Verification_F | CCAACCGCAGTAACGAATTG |
| IS5-KO_Verification_R | CCAGATCCTAATGTACCGTCTC |
| Aux3_KO_6kb_F | GCAGAGCAAGCATCACAG |
| Aux3_KO_6kb_R | ACCAGCTTTAATAACTGCTTG |
| lacZ-WT-6kb_pCVD442-SmaI_F | atgcgatatcgagctctcccgggACCCTAAGCGGTTCAATTTTG |
| lacZ-WT-6kb_pCVD442-SmaI_R | taacaatttgtggaattcccgggTGATCCGATGATCTTTTCG |
| **Allelic Exchange** | |
| *int*-Upstream_pCVD442-SmaI_F | accgcatgcgatatcgagctctcccgggCCTGCTGATAAAGCGGCG |
| *int*-Upstream_KanFRT_R | tccagcctacCTTGCATCAGAAAATTTGGTACAAATTTG |
| KanFRT_*int*-Upstream_F | ctgatgcaagGTAGGCTGGAGCTGCTTC |
| KanFRT_*int*-Downstream_R | tgatctcataATTCCGGGGATCCGTCGAC |
| *int*-Downstream_KanFRT_F | tccccggaatTATGAGATCATAGCAACCATC |
| *int*-Downstream_pCVD442-SmaI_R | gcggataacaatttgtggaattcccgggCGTCGTATCAGTTGGTCG |
| *int*-KO-Verification_F | AACTCGTTACGCCGCTATTT |
| *int*-KO-Verification_R | GTCACCTCATCCTTGCTGTT |
| KanFRT_*is5*-Downstream_R | ttggcggattATTCCGGGGATCCGTCGAC |
| *is5*-Downstream _KanFRT_F | tccccggaatAATCCGCCAATAGCCGGAG |
| *is5*-Downstream _pCVD442-SmaI_R | gcggataacaatttgtggaattcccgggTTGATGCTCTGTCCATCC |
| *int-is5*-KO-Verification_F | AACTCGTTACGCCGCTATTT |
| *int-is5*-KO-Verification_R | CCAGATCCTAATGTACCGTCTC |
| $int^{AM19226}$-Upstream_pCVD442-SmaI_F | accgcatgcgatatcgagctctcccgggAGTGCCTGCTGATAAAGC |
| $int^{AM19226}$-Upstream _KanFRT_R | gctccagcctacACTCACCTTGCATCAGAAAATTTG |
| KanFRT_$int^{AM19226}$-Upstream _F | atgcaaggtgagtGTAGGCTGGAGCTGCTTC |

| | |
|---|---|
| KanFRT_*int*$^{AM19226}$-Downstream_R | tatgtttatccatATTCCGGGGATCCGTCGAC |
| *int*$^{AM19226}$-Downstream_KanFRT_F | gatccccggaatATGGATAAACATAGTAATAGTGAC |
| *int*$^{AM19226}$-Downstream_pCVD442-SmaI_R | gcggataacaatttgtggaattcccgggTTAAATTTGAACAGTCAAAACTATAG |
| *int*$^{AM19226}$-KO_Verification_F | TGTAACTCACCAGTCACCTTTC |
| *int*$^{AM19226}$-KO_Verification_R | GACAACTCATTCCAGCCATAGA |
| **Transcomplementation** | |
| All-*int*-$P_{end}$_pGP704-mTn7-NotI_F | ggatccacgcgtcttaaggcTTTCTTCAGTGTCATTCTTATG |
| *int*$^{A1552}$_pGP704-mTn7-NotI_R | cccgacgggcccggtaccgcTCAGAGACTTAATCGCATC |
| *int*$^{AM19226}$_pGP704-mTn7-NotI_R | cccgacgggcccggtaccgcTTATCCATTATAATAAATTCCTAGAATAC |
| *int*$^{1154-74}$_pGP704-mTn7-NotI_R | cccgacgggcccggtaccgcTTAGAAAATAATACTTGTCCACTTC |
| FLAG-All-*int*_pGP704-mTn7-araC-$P_{BAD}$-NcoI_F | ggctagcaggaggaattcacc***atggactacaaagacgatgacgacaag***GCAATAACGGATGCATGG |
| *int*$^{A1552}$_pGP704-mTn7-araC-$P_{BAD}$-NcoI_R | tctagaggatccccgggtacTCAGAGACTTAATCGCATCTTC |
| *int*$^{AM19226}$_pGP704-mTn7-araC-$P_{BAD}$-NcoI_R | tctagaggatccccgggtacTTATCCATTATAATAAATTCCTAGAATACTC |
| **Donor Vectors** | |
| Aux3_Frag1_F | aagcgtatttTTTCACACAAGTGTTGCTTG |
| Aux3_Frag1_*attP*2_KO_F | cataaagcatAACTCACCAGTCACCTTTCATT |
| Aux3_Frag1_R | TGCCGATGAAGAAGATGTATG |
| Aux3_Frag2_F | AACAAGACAAGTCTGAACG |
| Aux3_Frag2_R | ttgtgtgaaaAAATACGCTTTAAATTCAATG |
| Aux3_Frag2_*attP*2_KO_R | ctggtgagttATGCTTTATGAATTTTGCG |
| *attP*_pKNOCK-Kan-SmaI_F | gctctagaactagtggatccCAGCAACGGGTCTCAGTATTG |
| *attP*_pKNOCK-Kan-SmaI_R | ttgatatcgaattcctgcagCTGTGGCTTATCTCAGTCTTAC |
| **Other** | |
| pUC19_Insert_F | GGAAACAGCTATGACCATGATTAC |
| pUC19_Insert_R | GGGTAACGCCAGGGTTT |
| pCVD442_insert_F | ACTAAATAATAGTGAACGGCAGGTA |
| pCVD442_insert_R | GTGAGCGGATAACAATTTGTGG |
| pBAD24_insert_F | GGCGTCACACTTTGCTATG |
| pBAD24_insert_R | GTTCTGATTTAATCTGTATCAGGCT |
| pKNOCK_ins_F | GGGATGTAACGCACTGAGAA |
| pKNOCK_ins_R | CGGATTCACCACTCCAAGAA |

* Gibson overlaps are shown as lowercase letters. When Gibson assembly involves three fragments, primers are named such that the first half of the name describes the amplified region and the second half describes the region to which the overlap is complementary.

**Table S3.** Genomes Used In This Study

| Organism | Strain | Serotype | Biotype | Ref Seq | Assembly Level |
|---|---|---|---|---|---|
| *V. cholerae* | N16961 | O1 | El Tor | GCF_000006745.1 | Complete |
| *V. cholerae* | A1552 | O1 | El Tor | GCF_002892855.1 | Complete |
| *V. cholerae* | M66-2 | O1 | El Tor | GCF_000021605.1 | Complete |
| *V. cholerae* | MAK 757 | O1 | El Tor | GCF_000153865.1 | Scaffold |
| *V. cholerae* | C6706 | O1 | El Tor | GCF_000237785.1 | Contig |
| *V. cholerae* | CIRS101 | O1 | El Tor | GCF_000175695.1 | Contig |
| *V. cholerae* | CP1041 | O1 | El Tor | GCF_000279245.1 | Contig |
| *V. cholerae* | MJ-1236 | O1 | El Tor | GCF_000022585.1 | Complete |
| *V. cholerae* | 2010EL-1786 | O1 | El Tor | GCF_000166455.1 | Complete |
| *V. cholerae* | HC-07A1 | O1 | El Tor | GCF_000318485.2 | Contig |
| *V. cholerae* | BX 330286 | O1 | El Tor | GCF_000174335.1 | Contig |
| *V. cholerae* | O395 | O1 | Classical | GCF_000016245.1 | Complete |
| *V. cholerae* | V52 | O37 | | GCF_000167935.2 | Scaffold |
| *V. cholerae* | MS6 | O1 | | CF_000829215.1 | Complete |
| *V. cholerae* | AM-19226 | O39 | | GCF_000153785.2 | Scaffold |
| *V. cholerae* | PA1849 | O1 | Classical | NA | |
| *V. cholerae* | CA401 | O1 | Classical | NA | |
| *V. cholerae* | A76 | O1 | Classical | GCF_001259495.1 | Scaffold |
| *V. cholerae* | A60 | O1 | Classical | GCF_001248195.1 | Scaffold |
| *V. cholerae* | A68 | O1 | Classical | GCF_001259635.1 | Scaffold |
| *V. cholerae* | A111 | O1 | Classical | GCA_001253495.1 (GenBank) | Scaffold |
| *V. cholerae* | A59 | O1 | Classical | GCF_001254535.1 | Scaffold |
| *V. cholerae* | A49 | O1 | Classical | GCA_001253835.1 (GenBank) | Scaffold |
| *V. cholerae* | A57 | O1 | Classical | GCA_001250255.1 (GenBank) | Scaffold |
| *V. cholerae* | A51 | O1 | Classical | GCA_001253435.1 (GenBank) | Scaffold |
| *V. cholerae* | A46 | O1 | Classical | GCF_001259555.1 | Scaffold |
| *V. cholerae* | NIH41 | O1 | Classical | GCF_000736865.1 | Contig |
| *V. cholerae* | RC27 | O1 | Classical | GCF_000176395.1 | Contig |
| *V. cholerae* | A279 | O1 | Classical | GCA_001253555.1 (GenBank) | Scaffold |
| *V. cholerae* | A61 | O1 | Classical | GCF_001250935.1 | Scaffold |
| *V. cholerae* | A50 | O1 | Classical | GCA_001254735.1 (GenBank) | Scaffold |
| *V. cholerae* | A103 | O1 | Classical | GCA_001254575.1 (GenBank) | Scaffold |
| *V. cholerae* | A70 | O1 | Classical | GCF_001248905.1 | Scaffold |
| *V. cholerae* | GP16 | O1 | Classical | GCF_001251495.1 | Scaffold |
| *V. cholerae* | GP8 | O1 | Classical | GCF_001253575.1 | Scaffold |
| *V. cholerae* | A389 | O1 | Classical | GCA_001259795.1 (GenBank) | Scaffold |
| *V. cholerae* | A66 | O1 | Classical | GCF_001260915.1 | Scaffold |
| *V. cholerae* | 2740-80 | O1 | | GCF_000168915.2 | Scaffold |
| *V. cholerae* | NCTC8457 | O1 | El Tor | GCF_000153945.1 | Scaffold |
| *V. cholerae* | HC38-A1 | O1 | El Tor | GCF_000221485.1 | Scaffold |
| *V. cholerae* | HC33-A2 | O1 | El Tor | GCF_000234885.1 | Scaffold |
| *V. cholerae* | HC32-A1 | O1 | El Tor | GCF_000234905.1 | Scaffold |
| *V. cholerae* | FC1225 | O139 | | GCF_002194335.1 | Contig |

| | | | | | |
|---|---|---|---|---|---|
| *V. cholerae* | FC2273 | O139 | | GCF_002194215.1 | Contig |
| *V. cholerae* | FC2271 | O139 | | GCF_002194235.1 | Contig |
| *V. cholerae* | FC1341 | O139 | | GCF_002194265.1 | Contig |
| *V. cholerae* | FC3611a | O139 | | GCF_002194165.1 | Contig |
| *V. cholerae* | FC1384 | O139 | | GCF_002194245.1 | Contig |
| *V. cholerae* | FC3611b | O139 | | GCF_002194185.1 | Contig |
| *V. cholerae* | FC1105 | O139 | | GCF_002194295.1 | Contig |
| *V. cholerae* | CP1041 | O1 | El Tor | GCF_000279245.1 | Contig |
| *V. cholerae* | MO10 | O139 | | GCF_000152425.1 | Scaffold |
| *V. cholerae* | FC1877 | O139 | | GCF_002194155.1 | Contig |
| *V. cholerae* | FC1817 | O139 | | GCF_002194305.1 | Contig |
| *V. cholerae* | IEC224 | O1 | El Tor | GCF_000250855.1 | Complete |
| *V. cholerae* | A6 | O1 | El Tor | GCF_001255575.1 | Scaffold |
| *V. cholerae* | 1154-74 | O49 | | GCF_000969235.1 | Complete |
| *V. cholerae* | 2479-86 | O1 | | GCF_001857305.1 | Contig |
| *V. cholerae* | 2012Env-9 | O1 | | GCF_000788715.2 | Complete |
| *V. cholerae* | Env-390 | O1 | | GCF_001854425.1 | Complete |
| *V. cholerae* | 20000 | nonO1/O139 | | GCF_004328575.1 | Complete |
| *V. cholerae* | P-18748 | nonO1/O139 | | GCF_002196055.1 | Contig |
| *V. cholerae* | HE-39 | nonO1/O139 | | GCF_000220765.2 | Contig |
| *V. cholerae* | HC-43B1 | O1 | | GCF_000279435.1 | Contig |
| *V. cholerae* | TM11079-80 | O1 | | GCF_000174255.1 | Contig |
| *V. cholerae* | HE-45 | nonO1/O139 | | GCF_000279285.1 | Contig |
| *V. cholerae* | 1587 | O12 | | GCF_000168895.2 | Scaffold |
| *V. cholerae* | 12129(1) | O1 | | GCF_000174115.1 | Contig |
| *V. cholerae* | TMA21 | nonO1/O139 | | GCF_000174295.1 | Contig |
| *V. cholerae* | DL4211 | O123 | | GCF_001953365.1 | Scaffold |
| *V. cholerae* | 623-39 | nonO1/O139 | | GCF_000154005.2 | Scaffold |
| *V. cholerae* | HE-25 | nonO1/O139 | | GCF_000279265.1 | Contig |
| *V. cholerae* | DL4215 | O113 | | GCF_001953375.1 | Scaffold |
| *V. cholerae* | MZO-2 | O14 | | GCF_000153985.2 | Scaffold |
| *V. cholerae* | MZO-3 | O37 | | GCF_000168935.2 | Scaffold |
| *V. cholerae* | 571-88 | O105 | | GCF_000736945.1 | Contig |
| *V. cholerae* | 234-93 | O141 | | GCF_000737005.1 | Contig |
| *V. cholerae* | 3568-07 | O141 | | GCF_001857505.1 | Contig |
| *V. cholerae* | V51 | O141 | | GCF_000152465.2 | Scaffold |
| *V. cholerae* | CP1110 | O75 | | GCF_000387585.1 | Contig |
| *V. cholerae* | CP1111 | O75 | | GCF_000387625.1 | Contig |
| *V. cholerae* | CP1112 | O75 | | GCF_000387645.1 | Contig |
| *V. cholerae* | CP1113 | O75 | | GCF_000387665.1 | Contig |
| *V. cholerae* | CP1114 | O75 | | GCF_000387685.1 | Contig |
| *V. cholerae* | CP1115 | O75 | | GCF_000387605.1 | Contig |
| *V. cholerae* | CP1116 | O75 | | GCF_000387725.1 | Scaffold |
| *V. cholerae* | CP1117 | O75 | | GCF_000387705.1 | Contig |
| *Vibrio* sp. | 2015V-1076 | | | GCF_003311815.1 | Contig |
| *Vibrio* sp. | 2017V-1038 | | | GCF_003311805.1 | Contig |
| *Vibrio* sp. | 2017V-1070 | | | GCF_003311865.1 | Contig |
| *Vibrio* sp. | 2016V-1062 | | | GCF_003311825.1 | Contig |
| *Vibrio* sp. | 2017V-1085 | | | GCF_003311895.1 | Contig |
| *Vibrio* sp. | 2523-88 | | | GCF_003311755.1 | Contig |
| *Vibrio* sp. | 2016V-1018 | | | GCF_003312035.1 | Contig |
| *Vibrio* sp. | 2017V-1124 | | | GCF_003311885.1 | Contig |
| *V. mimicus* | SX-4 | | | GCF_000222145.1 | Scaffold |

**Table S4.** Aux3 Enrichment Analysis

| | ***tseL/vasX/vgrG3* Grades ≥ 99%** | ***tseL/vasX/vgrG3* Grades < 99%** | **Total** |
|---|---|---|---|
| ***tseH* Grade ≥ 99%** | 461 | 1 | 462 |
| ***tseH* Grade < 99%** | 86 | 24 | 110 |
| **Total** | 547 | 25 | 572 |
| Fisher's Exact Test: $p = 2.2 \times 10^{-16}$ | | | |

**Table S5.** Aux3 Transfer Experiment Conjugative Frequencies

| Experiment | | Donor CFU/mL | Recipient CFU/mL | Transconjugant CFU/L | Conjugation Frequency |
|---|---|---|---|---|---|
| E.V. x Tn | A | 4.07E+08 ± 1.80E+08 | 1.03E+09 ± 3.00E+08 | 199.00 ± 0.00 | 2.08E-07 ± 7.32E-08 |
| | D | 2.73E+08 ± 6.43E+07 | 2.47E+09 ± 1.79E+09 | 199.00 ± 0.00 | 1.48E-07 ± 1.50E-07 |
| E.V. x Tn::*int*$^{P}$ | A | 2.13E+08 ± 9.24E+07 | 1.00E+09 ± 3.46E+08 | 199.00 ± 0.00 | 2.21E-07 ± 9.57E-08 |
| | D | 2.25E+08 ± 1.36E+08 | 1.01E+09 ± 5.06E+08 | 199.00 ± 0.00 | 2.53E-07 ± 1.73E-07 |
| E.V. x Tn::*int*$^{E}$ | A | 2.80E+08 ± 2.00E+07 | 3.32E+09 ± 4.07E+09 | 266.00 ± 116.05 | 2.22E-07 ± 1.74E-07 |
| | D | 3.47E+08 ± 1.21E+08 | 1.80E+09 ± 3.46E+08 | 199.33 ± 0.58 | 1.13E-07 ± 1.98E-08 |
| *attP*$^{WT}$ x Tn | A | 2.07E+08 ± 4.16E+07 | 5.67E+08 ± 2.77E+08 | 266.00 ± 116.05 | 5.25E-07 ± 2.28E-07 |
| | D | 2.33E+08 ± 1.17E+08 | 1.53E+09 ± 1.03E+09 | 800.00 ± 346.41 | 1.05E-06 ± 1.26E-06 |
| *attP*$^{WT}$ x Tn::*int*$^{P}$ | A | 2.20E+08 ± 7.21E+07 | 5.13E+08 ± 1.03E+08 | 666.67 ± 115.47 | 1.33E-06 ± 2.83E-07 |
| | D | 2.47E+08 ± 1.03E+08 | 2.27E+09 ± 1.42E+09 | 199.67 ± 0.58 | 1.17E-07 ± 7.47E-08 |
| *attP*$^{WT}$ x Tn::*int*$^{E}$ | A | 1.40E+08 ± 7.21E+07 | 1.13E+09 ± 3.16E+08 | 1.33E+05 ± 5.77E+04 | 1.22E-04 ± 4.79E-05 |
| | D | 1.47E+08 ± 1.01E+08 | 1.53E+09 ± 9.45E+08 | 733.00 ± 757.54 | 1.89E-07 ± 5.17E-08 |
| *attP*$^{KO}$ x Tn | A | 2.73E+08 ± 7.02E+07 | 8.00E+08 ± 4.00E+08 | 266.33 ± 115.76 | 3.60E-07 ± 1.26E-07 |
| | D | 2.40E+08 ± 1.59E+08 | 2.07E+09 ± 1.01E+09 | 199.33 ± 0.58 | 1.19E-07 ± 7.06E-08 |
| *attP*$^{KO}$ x Tn::*int*$^{P}$ | A | 2.47E+08 ± 5.77E+07 | 6.67E+08 ± 4.62E+08 | 399.67 ± 200.50 | 8.89E-07 ± 6.74E-07 |
| | D | 2.07E+08 ± 1.29E+08 | 2.67E+09 ± 1.90E+09 | 266.33 ± 115.76 | 1.37E-07 ± 9.66E-08 |
| *attP*$^{KO}$ x Tn::*int*$^{E}$ | A | 2.33E+08 ± 5.77E+07 | 8.67E+08 ± 5.03E+08 | 333.00 ± 231.23 | 4.64E-07 ± 3.06E-07 |
| | D | 1.57E+08 ± 1.45E+08 | 2.63E+09 ± 2.65E+09 | 199.33 ± 0.58 | 1.82E-07 ± 1.92E-07 |
| Tn only | A | 199.67 ± 0.58 | 7.67E+08 ± 3.79E+08 | 199.00 ± 0.00 | 2.99E-07 ± 1.20E-07 |
| | D | 999.67 ± 1216.88 | 1.53E+09 ± 7.02E+08 | 199.67 ± 0.58 | 2.05E-07 ± 1.70E-07 |
| Tn::*int*$^{P}$ only | A | 266.00 ± 116.05 | 9.20E+08 ± 4.85E+08 | 199.00 ± 0.00 | 2.95E-07 ± 2.23E-07 |
| | D | 266.00 ± 116.05 | 2.20E+09 ± 1.25E+09 | 199.00 ± 0.00 | 1.11E-07 ± 5.53E-08 |
| Tn::*int*$^{E}$ only | A | 199.00 ± 0.00 | 8.67E+08 ± 2.31E+08 | 199.00 ± 0.00 | 2.43E-07 ± 7.66E-08 |
| | D | 999.67 ± 1385.93 | 1.80E+09 ± 3.46E+08 | 199.00 ± 0.00 | 1.14E-07 ± 2.46E-08 |
